# Disentangling metacommunity processes using multiple metrics in space and time

**DOI:** 10.1101/2020.10.29.361303

**Authors:** Laura Melissa Guzman, Patrick L. Thompson, Duarte S Viana, Bram Vanschoenwinkel, Zsófia Horváth, Robert Ptacnik, Alienor Jeliazkov, Stéphanie Gascón, Pieter Lemmens, Maria Anton-Pardo, Silke Langenheder, Luc De Meester, Jonathan M. Chase

## Abstract

Metacommunity ecology has focused on using observational and analytical approaches to disentangle the role of critical assembly processes, such as dispersal limitation and environmental filtering. Many methods have been proposed for this purpose, most notably multivariate analyses of species abundance and its association with variation in spatial and environmental conditions. These approaches tend to focus on few emergent properties of metacommunities and have largely ignored temporal community dynamics. By doing so, these are limited in their ability to differentiate metacommunity dynamics. Here, we develop a Virtual ecologist’ approach to evaluate critical metacommunity assembly processes based on a number of summary statistics of community structure across space and time. Specifically, we first simulate metacommunities emphasizing three main processes that underlie metacommunity dynamics (density-independent responses to abiotic conditions, density-dependent biotic interactions, and dispersal). We then calculate a number of commonly used summary statistics of community structure in space and time, and use random forests to evaluate their utility for understanding the strength of these three processes. We found that: (i) time series are necessary to disentangle metacommunity processes, (ii) each of the three studied processes is distinguished with different descriptors, (iii) each summary statistic is differently sensitive to temporal and spatial sampling effort. Some of the most useful statistics include the coefficient of variation of abundances through time and metrics that incorporate variation in the relative abundances (evenness) of species. Surprisingly, we found that when we only used a single snapshot of community variation in space, the most commonly used approaches based on variation partitioning were largely uninformative regarding assembly processes, particularly, variation in dispersal. We conclude that a combination of methods and summary statistics will be necessary to understand the processes that underlie metacommunity assembly through space and time.

## Introduction

A perennial goal amongst ecologists is to be able to infer processes that influence the emergent ecological patterns of interest. For example, in metacommunity ecology—the study of sets of local communities linked by the movement of organisms— a large body of work has focused on using statistical analyses to understand the relative importance of underlying processes in structuring community assembly (overviewed in Logue et al. 2011, Soininen 2014, Leibold and Chase 2017, Ovaskainen et al. 2019). Most notably, these processes include species interactions and environmental filtering that have dominated ‘niche-based’ thinking (e.g., Tilman 1982, Chase and Leibold 2003), as well as aspects of stochasticity and dispersal limitation inherent to ‘neutral-based’ perspectives (e.g., Hubbell 2001).

Unfortunately, following initial promise, it has become clear that inference-based analyses can suffer from statistical biases (e.g. Gilbert & Bennett, 2010) and that analyses of metacommunities at a single point in time (snapshot) are often insufficient to differentiate among multiple ecological processes. For example, early interest in using the shape of the species abundance distribution (SAD) alone for differentiating neutral theory from niche-based alternatives (Hubbell 2001, Volkov et al. 2003, McGill 2003) quickly gave way to the realization that multiple processes could produce a similar SAD shape (Chave et al. 2002, Wilson et al. 2003, Chisholm and Pacala 2010). Likewise, neutral theory’s predicted species-area and distance-decay relationships are also readily predicted from other metacommunity models (Condit et al. 2002).

Later, emphasis for disentangling metacommunity processes from patterns shifted to multivariate analyses of species composition and its spatial variation (i.e., beta-diversity) and how that is associated with variation in spatial (S) and environmental conditions (E). For example, Cottenie (2005) used multivariate variation partitioning (Borcard et al. 1992) to link empirical patterns of observed community structure (specifically, the fractions of variation explained by S vs. E) to four classic metacommunity archetypes. The strength of the relationship between environmental features and community composition was assumed to indicate the relative importance of environmentally driven species sorting processes, while the strength of the relationship between spatial features and community composition was assumed to indicate the degree to which communities were structured by dispersal (see also e.g., Legendre 2008, Soininen 2014, 2016). While this approach became widely used, the explicit connection between the metacommunity theories and this variation partitioning approach is weak. For example, a number of features can reduce the fraction of composition explained, including unmeasured spatial and environmental variables, biotic interactions, and temporal changes in environmental conditions (Gilbert and Bennett 2010, Smith and Lundholm 2010, Tucker et al. 2016, Leibold and Chase 2017). Indeed, most syntheses of these patterns show only a small amount of the variation in most metacommunities is explained (Cottenie 2005, Soininen 2014).

As a result of the limitations of the original multivariate partitioning, metacommunity ecologists have continued to develop more refined analytical tools. For example, some of the issues with the variation partitioning approach have been dealt with by including latent variables to account for unmeasured environmental variables (Peres-Neto and Legendre 2010), by correcting spurious correlations caused by autocorrelated environmental variables (Clappe et al. 2018), by improving the fit of environmental responses using tree-based machine learning (Viana et al. 2019), and/or by accounting for species co-occurrence patterns in the context of joint species distribution models (JSDM) (e.g. Ovaskainen et al. 2017). Other approaches include the use of multiple diversity indices (sometimes including functional and phylogenetic information), selected according to simulation models with known underlying dynamics (virtual ecologist approach) to assess possible echoes of different processes in complex empirical patterns (Munkemuller et al. 2012, Ovaskainen et al. 2019).

The approaches used to analyse metacommunity structure so far have mostly relied on analysis of spatial pattern alone, without considering temporal dynamics, and are only able to explain a relatively small amount of the variation observed in real metacommunities (Jabot et al. 2020). Temporal dynamics clearly play a critical role in many classic metacommunity models, and without considering time, it can be impossible to discern among processes. For example, communities that arise from neutral processes can lead to quite similar patterns of spatial compositional turnover compared to those that arise from priority effects (the order and timing of species arrival in a community), but their patterns diverge when temporal dynamics are considered (Tucker et al. 2016, Leibold and Chase 2017).

Here, we use a pluralistic, process-based approach to get closer to the ultimate goal of deriving process from pattern in community assembly and metacommunity dynamics (Figure 1). We build on a recent framework developed by Thompson et al. (2020), which leaves behind the idea that real metacommunities can be understood by comparing them to a set of discrete archetypes bound by restrictive assumptions (e.g., species sorting vs. mass effects vs. neutral). Instead, this framework emphasizes how a broad continuum of metacommunity dynamics can arise via the interplay of three key processes (building on e.g., Loreau et al. 2003, Gravel et al. 2006, Vellend 2010, 2016, Fournier et al. 2017): 1) density-independent responses to abiotic conditions (i.e. the fundamental abiotic niche), 2) density-dependent biotic interactions (e.g. competition), and 3) dispersal. Stochasticity is incorporated as the probabilistic realization of the first three core processes (Shoemaker et al. 2020) (Figure 1 - arrows i and ii).

**Figure 1:**
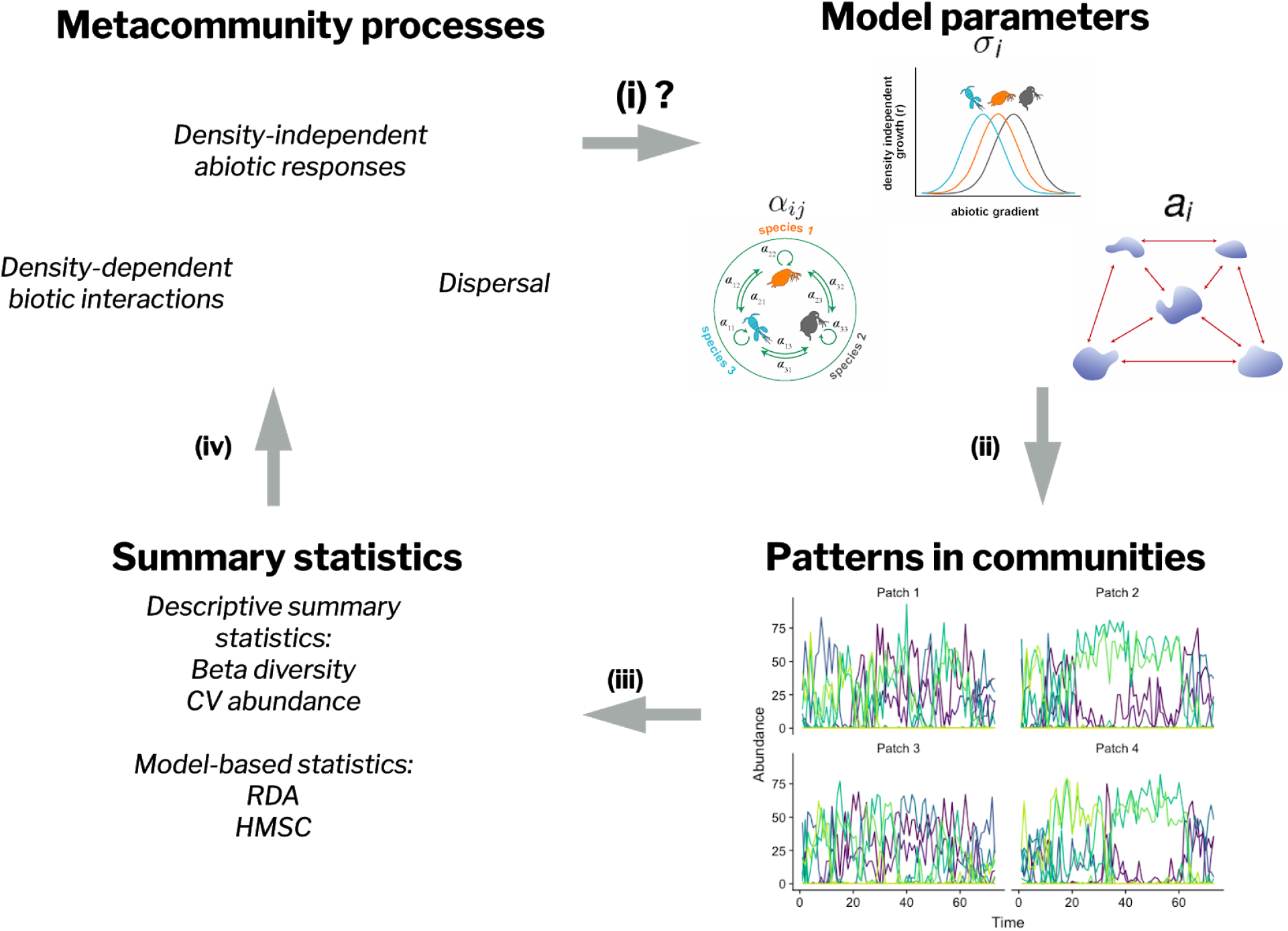
Workflow for finding summary statistics that distinguish patterns resulting from different metacommunity processes, (i) The different strengths of the processes result in different metacommunity dynamics, (ii) We adjusted key model parameters - abiotic niche breadth parameter (σ_*i*_), inter and intra competition strengths (*a*_ij_), and the probability of dispersal parameter(a_*i*_) - to change the strength of the density-independent, density-dependent, and dispersal processes in the simulations, (iii) We used descriptive statistics and model-based statistics to summarise the dynamics observed across the metacommunity through the time series (here we present some examples of summary statistics, for the full list see Table 1). (iv) Using random forests, we identified which of the summary statistics are most useful at distinguishing the three different model parameters.

**Table 1.**
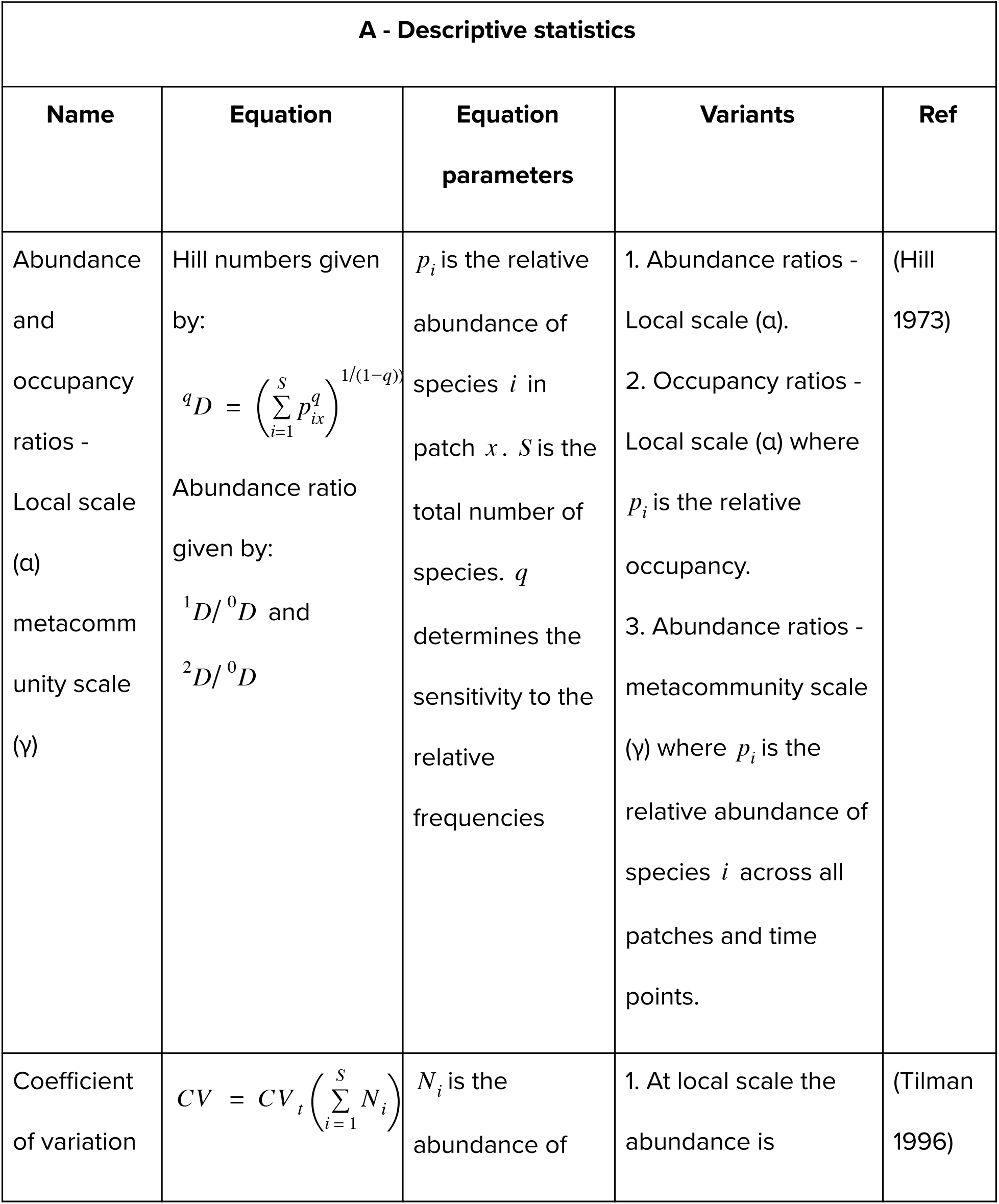
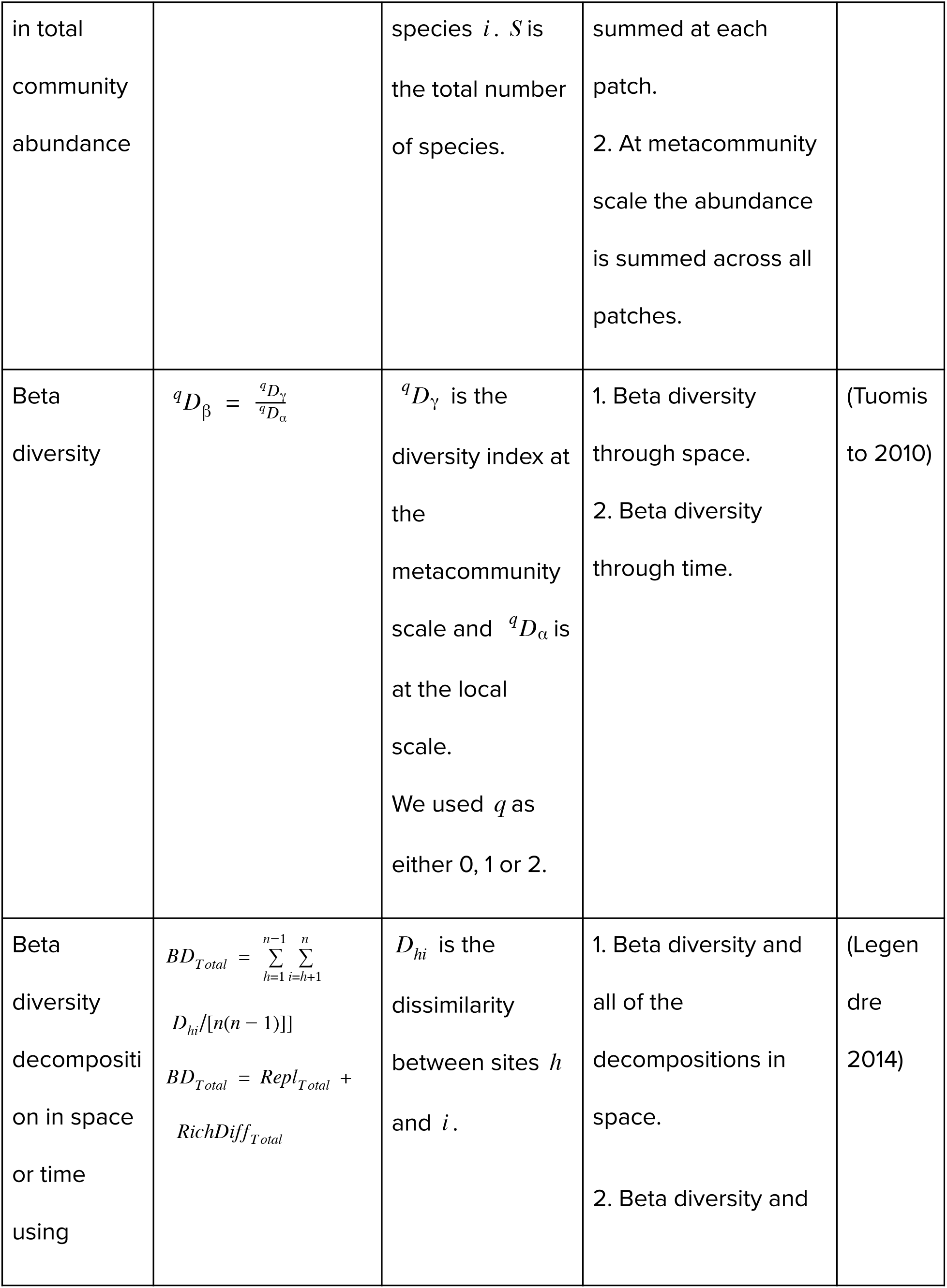
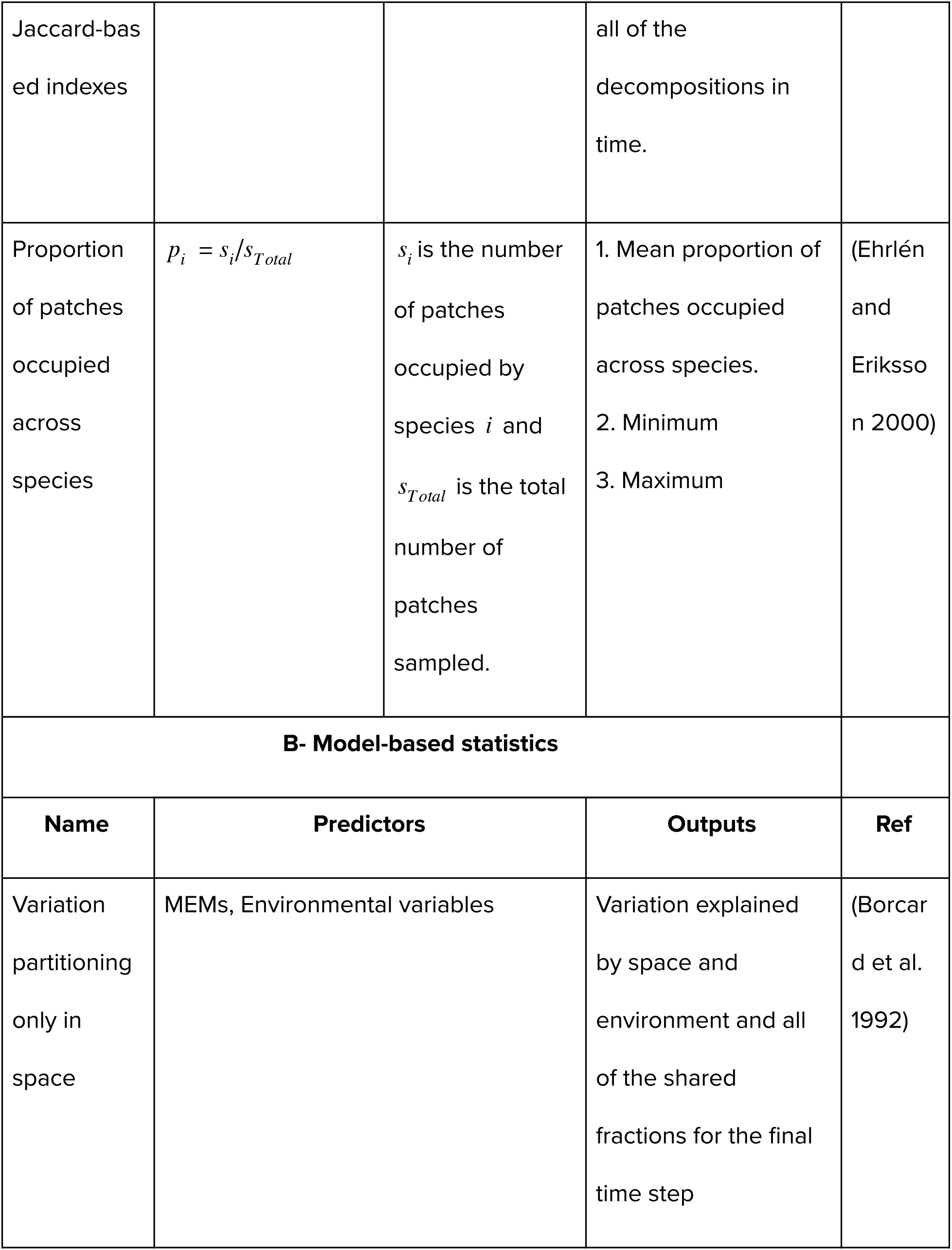
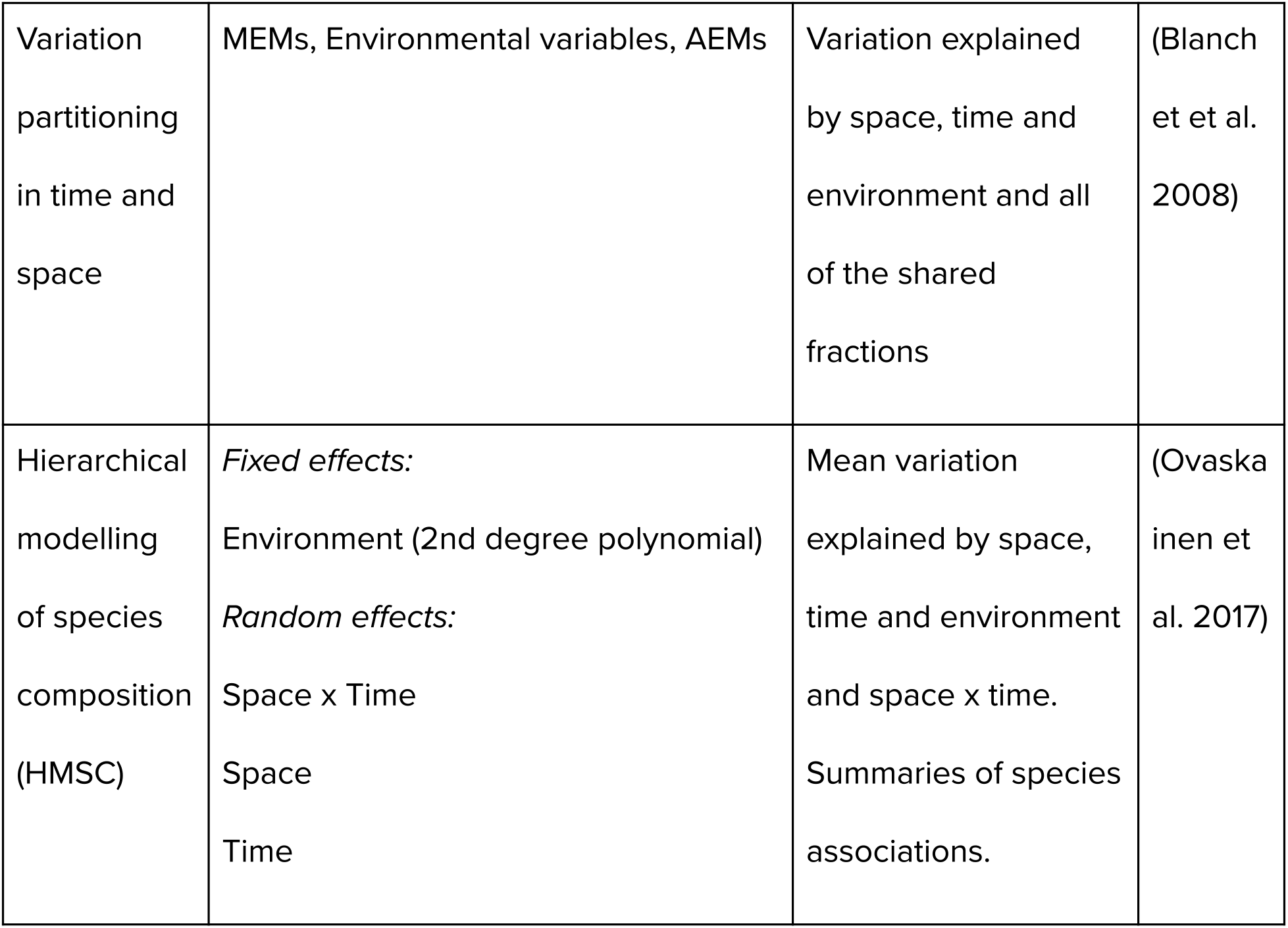
Main types of summary statistics investigated as candidate descriptors of metacommunity dynamics.

We use the model presented in Thompson et al. (2020), which simulates a wide range of metacommunity dynamics by changing the strength of each process. From simulated metacommunities, we estimate spatial and temporal variation in patterns of relative abundance, diversity and composition, to better capture the dynamic nature of metacommunity patterns (Figure 1 - arrow iii). We then use a suite of summary statistics - patterns and model-based statistics - in random forest models (non-parametric learning algorithms commonly used for regression and classification problems (Breiman 2001)) to predict the parameters of the simulation model that generated the metacommunity dynamics. This allowed us to identify the summary statistics that were most informative for distinguishing variation in each of the three underlying metacommunity processes (Figure 1 - arrow iv). Using these random forests we compared the performance of combinations of summary statistics and established the minimal set of summary statistics required to predict the metacommunity processes. As a final step, we examine the influence of sampling effort in time and space on the summary statistics.

Our results highlight that deriving pattern from process requires a pluralistic approach. That is, we find that there is no single ‘magic bullet’ parameter or analysis, but that a number of metacommunity statistics and analytical approaches may be required to assess the relative importance of the underlying processes. Finally we show which parameters are sensitive to data resolution by reducing the number of time points or patches, and conclude that most empirical studies may be severely undersampling their metacommunities to the point that reliable metrics cannot be obtained to infer metacommunity processes.

## Methods

### Simulations

Our goal with the simulations was to produce time series of metacommunity dynamics that varied in the three core processes in the model (abiotic niche breadth, density-dependent biotic interaction structure, and dispersal). Simulated metacommunity dynamics were generated using the model of Thompson et al. (2020) with replicate simulations (Model overviewed in Box 1). We used similar simulations presented in Thompson et al. (2020).

The metacommunities were composed of 100 patches and a starting regional richness of 50 species. We ran each simulation for 2200 time steps, including an initialization (200 time steps) and burn-in period (800 time steps), which were subsequently discarded. Following the burn-in period, we retained every 20^th^ time step to keep the size of the simulated data manageable, leaving a total of 60 retained time steps per simulation.

To change the density-independent responses to abiotic heterogeneity, we adjusted the abiotic niche breadth parameter (σ_*i*_) from a weak response to the environment (10) to a strong response to the environment (0.001). To change the density-dependent biotic responses, we adjusted the inter and intra competition strengths in four different scenarios (for details see Thompson et al. 2020): equal competition (α_*ij*_ = α_*ii*_), stabilizing competition (α_*ij*_ < α_*ii*_), mixed competition (α_*ij*_ can be less than or greater than α_*ii*_) and competition colonization tradeoff (30% of species are competitively dominant (α_*ij*_ > α_*ii*_) but their dispersal rates are an order of magnitude lower, 70% of species are subdominant and have stabilizing competitive interactions). Finally to change dispersal, we adjusted the probability of dispersal parameter (*a*_*i*_) ranging from metacommunities that were effectively disconnected (0.0001) to fully connected (0.464).

The abiotic environment in any given patch varies between 0 and 1 and is spatially and temporally autocorrelated. The mean value across time and space is 0.5.

We ran 660 combinations of dispersal rates, abiotic niche breadth and competition scenarios for 15 randomly generated replicate landscapes which varied in the location of the patches and the environmental conditions experienced. Landscapes were created by drawing xy coordinates in geographic space for each habitat patch from the range [1:100] and then converting these coordinates into a torus to avoid edge effects. Overall this resulted in 9900 simulation runs. Many of these combinations yielded no persistence in the metacommunity (2020 simulation runs) (Thompson et al. 2020). We define persistence as abundance greater than 0 after the initial burn in time. Lack of persistence mostly occurred when dispersal was low and responses to environmental variation were strong. We only considered simulations where the entire metacommunity (i.e., gamma diversity) had more than one species, and where the number of replicates that consistently showed persistence was greater than 10 out of 15. This ensured that parameter combinations that were not well replicated did not drive the patterns when we compared across parameter estimates.

### Summary statistics

We calculated 85 summary statistics from each simulation run. These included descriptive statistics (e.g, diversity metrics) as well as model-based statistics from commonly used methods to analyse metacommunity dynamics (e.g., variation partitioning fractions) (see Table 1 for the main description of the statistics and Table S1 for a full tally of all the statistics we used).

First, we calculated statistics that describe patterns of relative abundances (i.e., relative species commonness and rarity) that determine the shape of the SAD, including Hill number ratios based on either abundance or occupancy. These were based on the ratio of the Hill numbers, which reflect a measure of evenness while controlling for species richness (Hill 1973, Chao et al. 2014). We use two main ratios ^l^*D*/ ^0^*D* and ^2^*D*/ ^0^*D* to capture evenness. ^2^*D* is the inverse Simpson’s diversity, a measure of dominance in the community. ^1^*D* is the exponent of Shannon’s diversity and ^0^*D* is species richness. When a community is exactly even, ^2^*D* or ^1^*D* approximates ^0^*D* and therefore the ratio of these values is closer to 1. However, when the community is very uneven, then ^2^*D* or ^1^*D* will be much lower than ^0^*D* and the ratios of ^1^*D*/ ^0^*D* and ^2^*D*/ ^0^*D* will be much smaller.

Second, we calculated the coefficient of variation of local and regional abundance across all species through time. This statistic measures the stability of communities by providing a standardized index of variation in abundance through time (Tilman 1996, Loreau et al. 2003).

Third, we calculated beta diversity and its decomposition into richness differences and replacement components (Podani and Schmera 2011). Beta diversity measures compositional heterogeneity between habitat patches (spatial beta diversity) or time points (temporal beta diversity) (Tuomisto 2010), and it can be partitioned into species replacement and richness differences. Species replacement indicates the turnover of species among samples, for example due to environmental filtering or competition (Legendre 2014). On the other hand, richness differences may reflect the different coexistence parameters in different locations and/or dispersal limitation independently of species replacement (Schmera et al. 2020). Nestedness, for example, is a type of richness difference characterized by subsets of species from the richer site (Legendre 2014). Both species replacement and richness differences can be evaluated through time or through space.

Finally, we calculated the proportion of patches occupied for each species, and then calculated the mean, minimum, and maximum across species. Low occupancy can be a sign of dispersal limitation or the strength of competition, whereas high occupancy could point to mass effects (Ehrlén and Eriksson 2000).

For the model-based statistics, we used two families of statistical models to determine how the amount of variation in species’ abundances explained by time, space and environment are related to the processes that shape metacommunity structure. First, we performed a classical variation partitioning based on redundancy analysis (RDA) (Borcard et al. 1992) to quantify the amount of variation in community composition explained by the environment, space and time. We did this for data taken only at the final time point, using environment and space as predictors, and for data taken across the entire time series, using environment, space and time as predictors. For both models, we calculated the spatial component using distance-based Moran’s eigenvector maps (MEMs) calculated using the *dbmem* function from the *adespatial* package (Dray et al. 2019). The MEMs were calculated across all patches and the same set reused for all the simulations using the same landscape. We used all positively autocorrelated MEMs in the analysis, as selecting specific MEMs does not fully account for spatial autocorrelation in the residuals (Peres-Neto and Legendre 2010). While we recognize that using a selection procedure of MEMs can yield better results for individual simulation runs, we decided to keep the number of MEMs consistent across simulation runs so they are more easily comparable. We calculated the temporal component using asymmetric eigenvector maps (AEMs) calculated using the *aem.time* function from the *adespatial* package (Blanchet et al. 2008, Dray et al. 2019).

Second, we used Hierarchical Modelling of Species Communities (HMSC) (Ovaskainen et al. 2017), which is a hierarchical Bayesian joint species distribution model that uses fixed environmental predictors and spatiotemporal random effects to make community-level inference of assembly processes. We used species abundance with an assumed Poisson distribution as the response variable. The environmental variable was included as a fixed linear and quadratic effect to fit the Gaussian-shaped response of species to the abiotic environment. The spatial and temporal structures of the data were modelled through autocorrelated spatial, temporal, and spatiotemporal random factors. The spatial random effect was modelled using the x and y coordinates. The temporal random effect was modelled using timestep as a random temporal coordinate. The spatiotemporal random effect was modelled using the x, y, and time coordinates. We used the nearest-neighbour Gaussian process with 10 neighbours for the spatial and spatiotemporal random effects to reduce computation time (Tikhonov et al. 2020). We also restricted each random effect to a single latent factor to make the analysis computationally feasible. Although additional variation may be explained by allowing for more latent factors, it is unlikely to be large as there are no unmeasured environmental variables in our simulation model. The summary statistics we used in the Random Forest consisted of partitioned explained variation according to all fixed and random effects: environment, space, time, spacetime. We used both the raw variation fractions and standardized fractions by total TjurR^2^ to also account for differences in the amount of residual variation across simulations. In addition to the variation fractions, we used the estimates of species associations aggregated into statistics such as the proportion of positive or negative associations (Ovaskainen et al. 2019). HMSC was run across 4 chains, each with 1000 samples and a transient period of 5000 steps. Sensitivity analyses suggest that while this length of MCMC sampling does not lead to full convergence, it is sufficient to provide estimates of the summary statistics that we use in our analysis (Figure S7). Furthermore, restricting our analysis in this way was necessary to make it computationally feasible (following Ovaskainen et al. 2019). Thus, our results for the HMSC analysis should be comparable across the range of parameters in our simulations, but are conservative for the potential performance of HMSC in assessing metacommunity processes. HMSC was implemented using the HMSC-R package (Tikhonov et al. 2019).

##### Box 1 - Model

The dynamics of the metacommunity are governed by a Beverton-Holt (Beverton and Holt 1957) growth dynamics with generalized Lotka-Volterra competition:

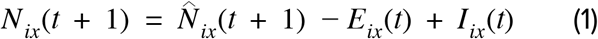

where 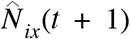 is the population size at time *t* before accounting for dispersal.

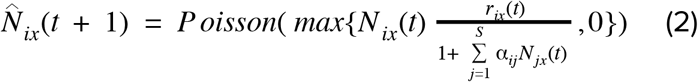

where 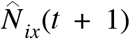 is the expected abundance of species *i* in patch *x* at time *t* + 1, α_*ij*_ is the per capita interaction effect of species *j* on species *i*, and *S* is the total number of species. Stochasticity in local demographic outcomes is incorporated through the Poisson draw in equation 2. *r*_*ix*_(*t*) is the density-independent growth rate of species *i* in patch *x* at time *t* which is given by:

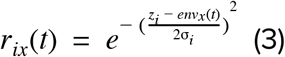

where *z*_*i*_ is the environmental optimum of species *i*, *env*_*x*_(*t*) is the environmental condition in patch *x* at time *t,* and σ_*i*_ is the abiotic niche breadth.

The number of emigrants *E*_*ix*_(*t*) was determined as the outcome of *N*_*ix*_(*t*) draws of a binomial distribution, each with a probability equal to *a*_*i*_. The destination of each of these emigrants is determined through a random draw of the patches, with their probabilities determined by:

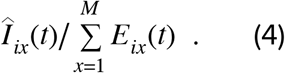

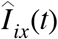 is the expected number of individuals that immigrate from species *i* in patch *x* at time *t* and it is given by:

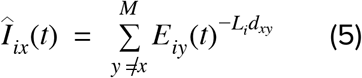

where *M* is the total number of patches, *E*_*iy*_(*t*) is the number of immigrants of species *i* from another patch *y*, *L*_*i*_ is the strength of exponential decrease of dispersal with distance and *d*_*xy*_ is the distance between patches *x* and *y*.

### Random Forests

We identified the summary statistics that were the most effective in discriminating among the metacommunity processes using random forests (Breiman 2001). We used random forests because they can be used both for regression and classification problems (dispersal and abiotic niche breadth parameters are continuous, while the competition scenario is a categorical parameter). Further, random forests are non-parametric, allowing any relationship between predictor and response, and can be used to rank the importance of the predictors (Breiman 2001).

We built 21 random forest models (7 classes of models for 3 predictors separately see Table 2) for each process separately to determine which summary statistics were better at predicting each process across all simulations (Table 2). Across all models, we ran 500 trees for each random forest. In random forests, the variable importance is determined by observing how much the prediction error increases when each variable is permuted, while the other variables remain the same (Breiman 2001, Liaw and Wiener 2002).

**Table 2.**
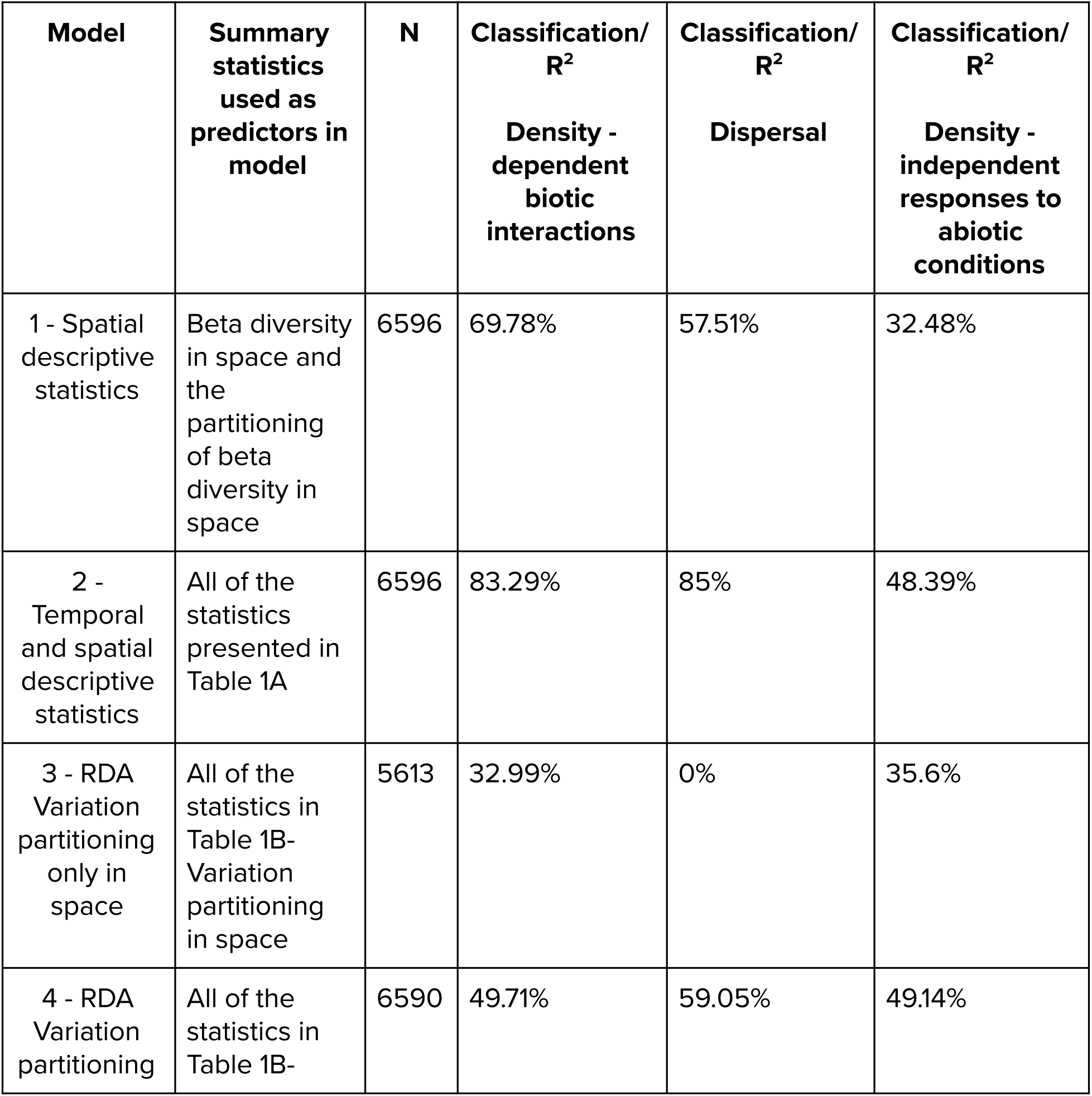
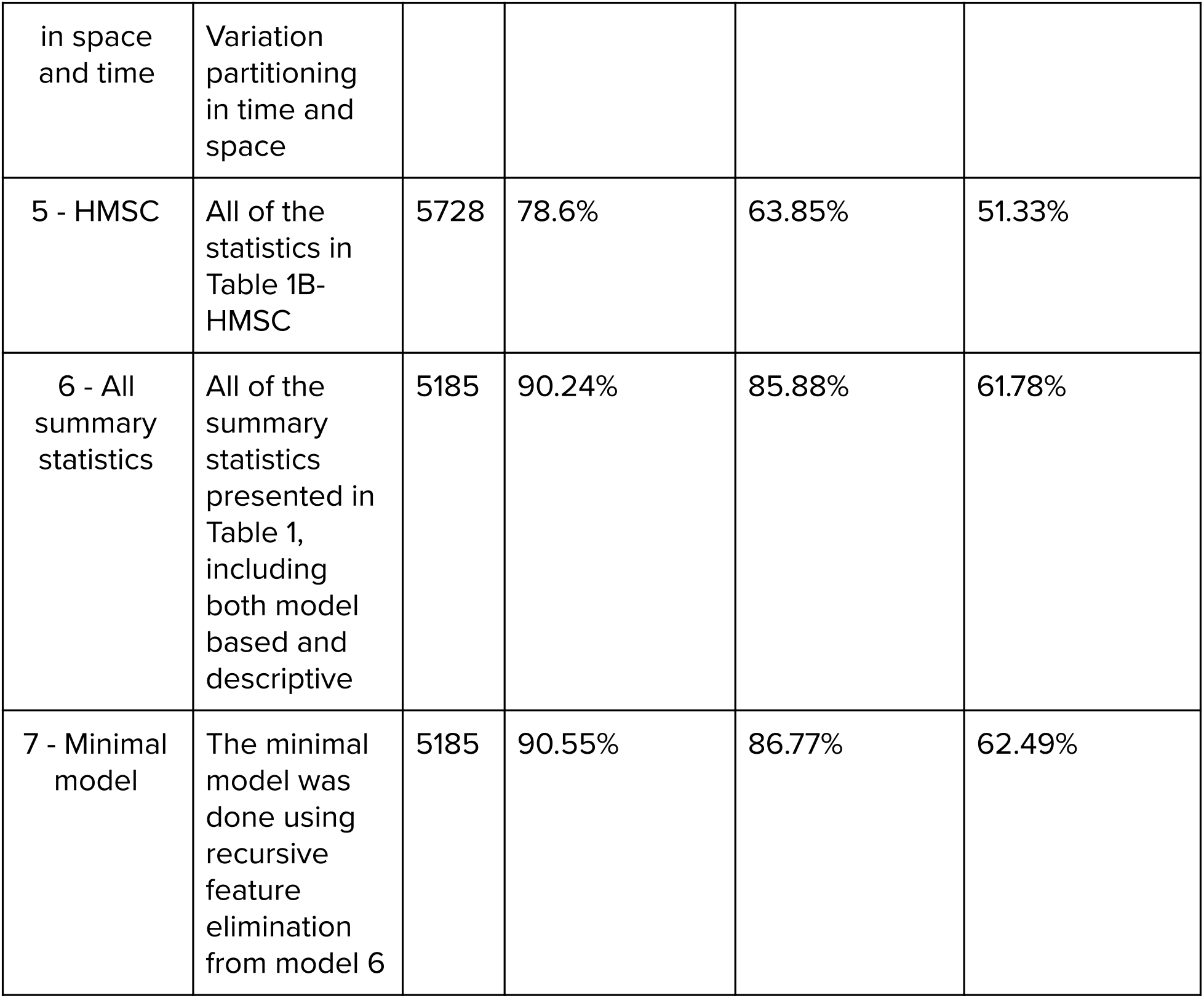
Random forests were used to compare the performance of sets of summary statistics at explaining variance in each of the three metacommunity processes: density-dependent biotic interactions, dispersal, density-independent responses to abiotic conditions. We set up 7 different types of models depending on the types of summary statistics for each of the three processes for a total of 21 random forests.

Because some simulations were discarded (see above), the maximum number of simulations we analyzed was 6596 (Table 2) for the simple metacommunity descriptors. For metacommunity descriptors derived from models (i.e. RDA or HMSC), we only ran random forests for metacommunities that had persistence of at least two species for more than 3 time points. In addition, some of the model-based statistics yielded NA values (e.g. when calculating species covariance involving transient species that occurred in only one time step). We removed the simulation results that had missing values. As a result, the number of simulations used in the random forest for the HMSC and all summary statistics was lower (5728 and 5185, respectively).

The minimal model was selected using recursive feature elimination and a 10-fold cross-validation procedure. This algorithm first partitions the data into test and training sets. The model is fit with all of the predictors to the training data and t tested with the held-back samples, where each predictor gets an importance value. The algorithm then keeps only the ‘n’ most important variables, re-fits the model and tests it again with the held-back samples. This procedure is repeated for 10 to 50 ‘n’ number of predictors (Figure S1). The algorithm determines the best number of predictors and the best predictors based on the prediction accuracy (with the held-back samples). The whole procedure is repeated 10 times (Ambroise and McLachlan 2002, Svetnik et al. 2004).

### Sensitivity analysis to spatial and temporal sampling effort

The summary statistics, with the exception of the spatial-only variation partitioning, were calculated on the entire simulation time series (after burn-in) of the metacommunity. However, empirical data are inevitably limited in the number of samples that can be obtained in space and time. Therefore, we investigated the effect of limited sampling by running our analyses on a subset of all existing patches and time points, as is typically the case in empirical studies. For this, we used a simulation that had stabilizing competition (α_*ij*_ < α_*ii*_), intermediate levels of density-independent responses to abiotic conditions (4.64159) and intermediate levels of dispersal (0.00215). We chose these parameter values because they yielded gamma diversity above 1 for all simulations and they were intermediate values of dispersal and abiotic conditions. Then, we randomly sampled m of 100 patches where m = 4, 8,12,16,…, 100. This sampling was repeated 1000 times to cover spatial variation in the subsamples. We also sampled t of our 60 time points, where t = 4, 8,12,16,…, 60. This sampling was not repeated, as the number of time points included was sequential. We subsampled time and space factorially (i.e., few time points and few patches, few time points and many patches, many time points and few patches, and many patches and time points).

All of the code for our simulations and analyses can be found on GitHub: https://aithub.com/lmauzman/disentanalinametacommunities, and will be mirrored at Zenodo upon acceptance.

## Results and Discussion

Overall, we found that jointly addressing time and space is necessary for distinguishing metacommunity dynamics. Including statistics that are measured through time increased the explanatory power of the random forests by up to 59% when compared to cases where only spatial variation was considered. This was the case when temporal variation was incorporated in the descriptive statistics (Table 2 - Model 1 vs Model 2) and in the RDA variation partitioning (Table 2 - Model 3 vs Model 4). These results suggest that a simple snapshot of communities at a certain moment in time is not sufficient, and neither substituting space for time in observational studies to understand spatio-temporal dynamics. Additionally, we found that different summary statistics are complementary and can capture different aspects of metacommunity dynamics. For instance, including both descriptive and model-based summary statistics increased the explanatory power of the random forests by up to 22% (Table 2 - Model 6 and 7).

For all processes, descriptive statistics were more informative than model-based statistics, even with fewer predictors (Table 2 - Model 2 vs Model 5). Model 2 included statistics that described communities through space *and* time. These statistics capture more variation in metacommunity dynamics than redundancy analysis or HMSC. For example, the variation of community biomass through time and the ratios of occupancy (evenness) are important statistics in the random forest (Figure 2). These statistics capture a real pattern in metacommunity variation, while variation partitioning (either RDA or HMSC) represent a model fit. Measures of temporal variability and the SADs are also important at distinguishing metacommunity processes. However, we emphasize that rather than using them in isolation, as had been the focus previously (e.g., distinguishing niche vs. neutral processes (Hubbell 2001, Volkov et al. 2003, McGill 2003, Chisholm and Pacala 2010), these statistics are much more informative when combined with other statistics such as beta diversity metrics (Table S1).

**Figure 2:**
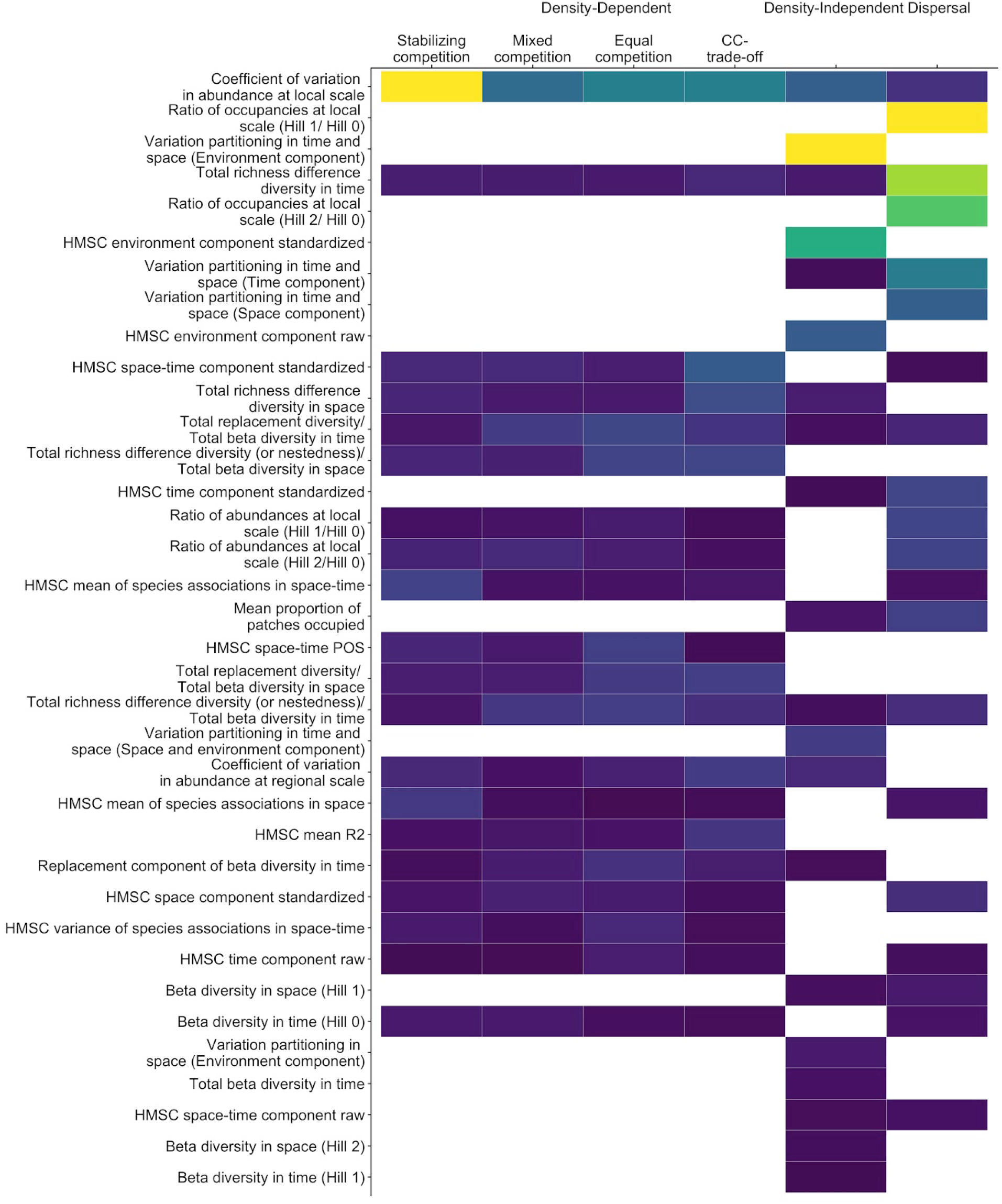
The best 20 performing summary statistics ordered from top to bottom by their overall importance in the minimal random forest model (model 7). The importance of the summary statistics decreases from top to bottom of the graph and from yellow (most important) to purple (least important). Blank areas are variables that were not selected in the minimal models. The most important summary statistic to differentiate between the four types of local interactions is the coefficient of variation in abundance at a local scale, the most important summary statistic for dispersal is the portion of total richness difference of beta diversity in time, and the most important summary statistic for the density-independent responses to abiotic conditions is the environmental component of the variation partitioning in time.

Despite its continued historic popularity, RDA based variation partitioning using only spatial data (Model 3) had the lowest performance to distinguish metacommunity dynamics (Table 2). This is likely because variation partitioning does not take into account species interactions and partly due to model misspecification (Viana et al. 2019). In addition, using a single snapshot in time does not capture the spatio-temporal dynamics of dispersal. While it is rather surprising that the random forest that included only spatial descriptive statistics performed better than the RDA-based variation partitioning in space, using metrics that describe community dynamics themselves rather than model-fits seems to be more informative for the random forest. Thus, despite the fact that this has been a very popular approach to infer potential metacommunity mechanisms, our analysis shows that this approach is prone to yield misleading interpretations.

Once we included temporal dynamics, the RDA based variation partitioning performed better in classifying and explaining the variation of the underlying processes (Table 2 - Model 4). Including time in the variation partitioning model increased the classification success for different types of biotic interactions to 50%, and explained more variation in dispersal (59%) and in responses to abiotic conditions (49%). It is especially remarkable that the amount of variation explained in dispersal was particularly sensitive to the inclusion of time, as it increased from 0% to 59% when time was included. Dispersal is a spatio-temporal process that affects the dynamics of local populations. Thus, it does not seem that surprising that including the effect of time, instead of the commonly used “snapshot” of the metacommunity, increases our predictive ability. Likewise, the random forest for biotic interactions and abiotic responses are more successful in inferring processes once we include time. As such, it seems clear that including sampling in both time and space is necessary for disentangling metacommunity processes, and that relying on snapshots in time for disentangling dynamic systems is insufficient.

Hierarchical modelling of species composition (HMSC) had a greater predictive power of metacommunity processes than redundancy analysis with time (Table 2 - Model 5 vs Model 4). The best improvement was in the classification of the biotic interactions (success rate 79% for model 5 vs. 50% for model 4). This is because the HMSC approach explicitly estimates positive or negative interspecific associations via species covariance *after* accounting for the effect of (environmental) covariates (Ovaskainen et al. 2017). Although the utility of using species associations for inferring the importance of actual species interactions is still widely debated (Blanchet et al. 2020), our results nevertheless suggest that these model based statistics increase our ability to distinguish the type of biotic interactions, even if the pairwise species interaction coefficients cannot be reliably recovered.

All statistics together, including descriptive and model-based statistics (Model 6), performed better for all three processes than using only one type of statistic alone. Several statistics provided complementary information that was useful at discriminating metacommunity processes (Table 2). However, this random forest included up to 85 predictors (All predictors in Table S1) and very little predictive ability was lost when we reduced it to a minimal model through backwards selection (Table 2). These minimal models included a subset of statistics depending on the random forest used to infer each of the processes (Figure 2). The best number of predictors was 50 for the density-dependent biotic interactions, 32 for dispersal and, 29 for density-independent responses to abiotic conditions (Figure S1). The most informative statistics selected by the random forest were those having the smallest variance at each parameter value, but exhibited most variation across parameter space (Figure 3a,e,i vs Figure 3c,d,h). In what follows, we describe the minimal model for each process.

**Figure 3:**
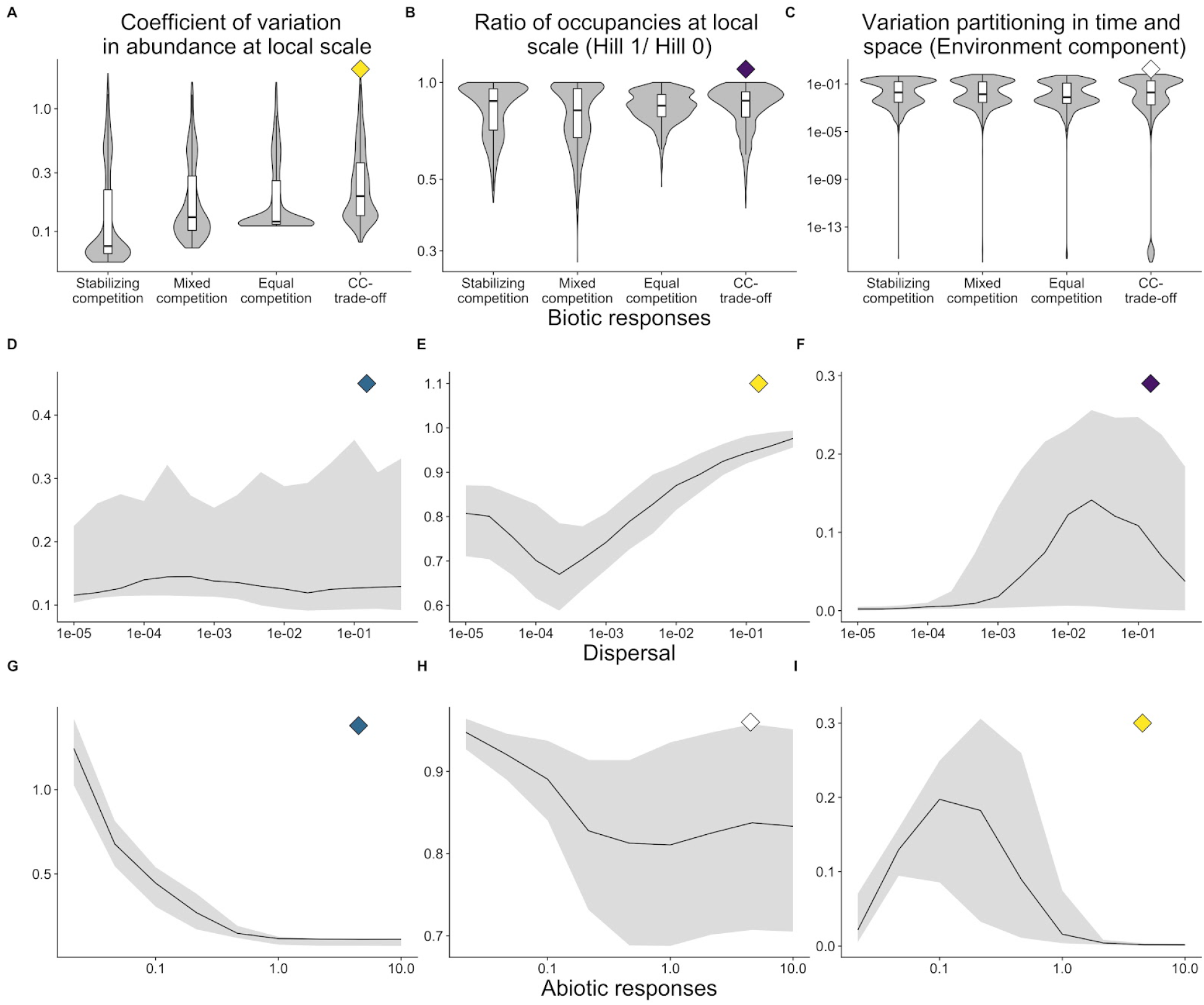
The three most important summary statistics for predicting the underlying metacommunity processes (density-dependent biotic interactions - top row; dispersal - middle row; and density-independent responses to abiotic conditions - bottom row) as determined by the minimal random forest model (model 6): 1) the coefficient of local scale variation in community abundance (left column), 2) ratio of occurrence at local scale (Hill 1/ Hill 0) (middle column), and 3) the environmental component of variation partition through time and space; right column. The color of the diamonds in the top right of each panel corresponds to the colour in Figure 2 and shows the relative importance of each summary statistic for explaining that metacommunity process. The error lines in the bottom two rows represent the 1^st^ and 3^rd^ quartiles of the values of the summary statistic, while the lines represent the median values of the summary statistic. The top row represents the distribution of the data using violins and boxplots within those violins represent the 1^st^ and 3^rd^ quartiles and the median of the summary statistics.

### Density-dependent biotic interactions

The minimal model to distinguish different types of biotic interactions had a classification success of 90.68%. Stabilizing competition was easiest to distinguish with the available summary statistics (3.32% error rate), while the competition-colonization trade-off was the hardest to separate from the other types (19.97% error rate). The most important summary statistic that helped to distinguish biotic interactions was the coefficient of variation of abundance at the local scale (Figures 2 and S2). This was lowest under stabilizing competition, increased with equal competition and mixed competition, and was highest with competition-colonization trade offs (Figure 3a). Under stabilizing competition, local diversity is higher (see also Thompson et al. 2020), which stabilizes overall community abundance through insurance and portfolio effects (Doak et al. 1998, Yatchi and Loreau 1999). The increased variability in the other scenarios is due to the stronger competitive effects (Chesson 2000), which reduce diversity, and can result in abrupt compositional transitions with greater temporal variability in community abundance (Thompson et al. 2020).

The second most important summary statistic was the proportion of spatial beta diversity due to richness differences. We found the lowest richness differences under equal competition and stabilizing competition, moderate richness differences with mixed competition and highest richness differences with competition-colonization trade offs. With stabilizing and equal competition, richness differences accounted for a smaller proportion of the total spatial beta diversity, meaning that patches often have a similar number of species and species replace each other across the landscape. On the other hand, with competition-colonization trade-offs, richness differences accounted for a higher proportion of spatial beta diversity (Thompson et al. 2020).

### Dispersal

The minimal model explained 86.77% of the variation in dispersal in the simulated metacommunities (Table 2). Here, community evenness ‒captured by the ^1^*D*/ ^0^*D* occupancy ratio— emerged as the most important summary statistic (Figure 2 and S3). This statistic has a U-shaped relationship with dispersal (Figure 3e). Low dispersal results in a relatively high evenness, which suggests that most species occupy a relatively similar number of patches, despite considerable variation in occupancy. At intermediate levels of dispersal, species mostly track environmental conditions, i.e. we find most effective species sorting. High dispersal again results in a relatively high evenness, suggesting that most species are distributed evenly across the landscape, pointing at mass effects (Figure 3e). The second important summary statistic was the total richness difference between time points (Figure 2 and S3, Table 1). We observed another U-shaped relationship with dispersal. When dispersal is low, species are largely constrained to their initial patches with little ability to move and are more likely to go extinct when environmental conditions change, leading to higher richness differences through time. As dispersal increases, species are maintained locally and less likely to go extinct, and therefore the richness differences between time points decreases (Leibold et al. 2004).

### Density-independent responses to abiotic conditions

The minimal model to explain variation in the strength of density independent environmental filtering in the simulated metacommunites captured only 62.5% of the variation, even with all the summary statistics available. The most important summary statistic for the responses to abiotic conditions was the environmental component of the variation partitioning through time and space (Figure 2 and S4). This component has a hump-shaped relationship with the strength of abiotic conditions (Figure 3i) and this relationship may be biological or statistical in nature. When species respond strongly to abiotic conditions (i.e. σ_*i*_ - niche breath- is small), the environmental component explains very little variation in community composition, because species will not be able to persist in their patches when the environment changes. As the responses to the abiotic conditions weaken (i.e. intermediate σ_*i*_), the environmental component explains the most variation in community composition. Under such conditions, species can respond to environmental variation by moving to suitable patches and not driven extinct by changes in environmental conditions before moving. Finally, when the responses to abiotic conditions are very weak (i.e. σ_*i*_ is large), the amount of variation explained by the environmental component is again low, as variation in environmental conditions does not lead to changes in community composition (Figure 3i). In addition, when niche breadth is very narrow (i.e. strong abiotic response), the models used here (both the E component generated via partial RDA or HMSC) may not be able to fit the environmental response (eg. RDA fits linear responses), but when the niche is wider, then the models are better able to fit the species response to the environment (either fitting the unimodal relationship like in HMSC or linear responses in RDA are more suitable). When the niche breath is too wide, environmental variation does not explain variation in species composition.

### Sensitivity analysis

When we reduced sample completeness to be more realistic for empirical studies, the summary statistics were not equally affected (Figure 4). We evaluated the error half-life of the summary statistics when time was fully sampled but patches were not, and vice-versa. The error half life is the minimum number of patches or sequential time points needed to reduce the ‘error’ in the summary statistic by half. Not surprisingly, we found that some summary statistics are more sensitive to the loss of patches while some summary statistics are more sensitive to the reduced coverage in time. Here, we describe the error half life for the number of patches when time is fully sampled, which is clear in the simulations, but less clear in empirical studies. Empiricists will have to determine what full sampling time means for their study system depending on the organisms studied. When time is fully sampled, total beta diversity in time, beta diversity in space (Hill 1 and 2), replacement diversity in time and richness differences in time, as well as all of the temporal and environmental components of variation partitioning, are robust to a reduced number of sampled patches. These statistics needed less than 8% of patches sampled to reduce the error rate by half. On the other hand, the spatial component of variation partitioning is very sensitive to the loss of patches, and reaches half the error at *72%* of patches remaining (Table S2, Figure S5). When space is fully sampled, the minimum and maximum proportion of patches occupied, the space-time component of variation partitioning, beta diversity in space and time (Hill 1 and 2), and the coefficient of variation in abundance at the local scale are very robust to the loss of time points, where only 8% of the time points are needed to reduce the error rate by half. On the other hand, the space-environment shared component of variation partitioning, and replacement and richness differences through time need more than 80% of time points to reduce the error rate in half (Table S3, Figure S6).

**Figure 4:**
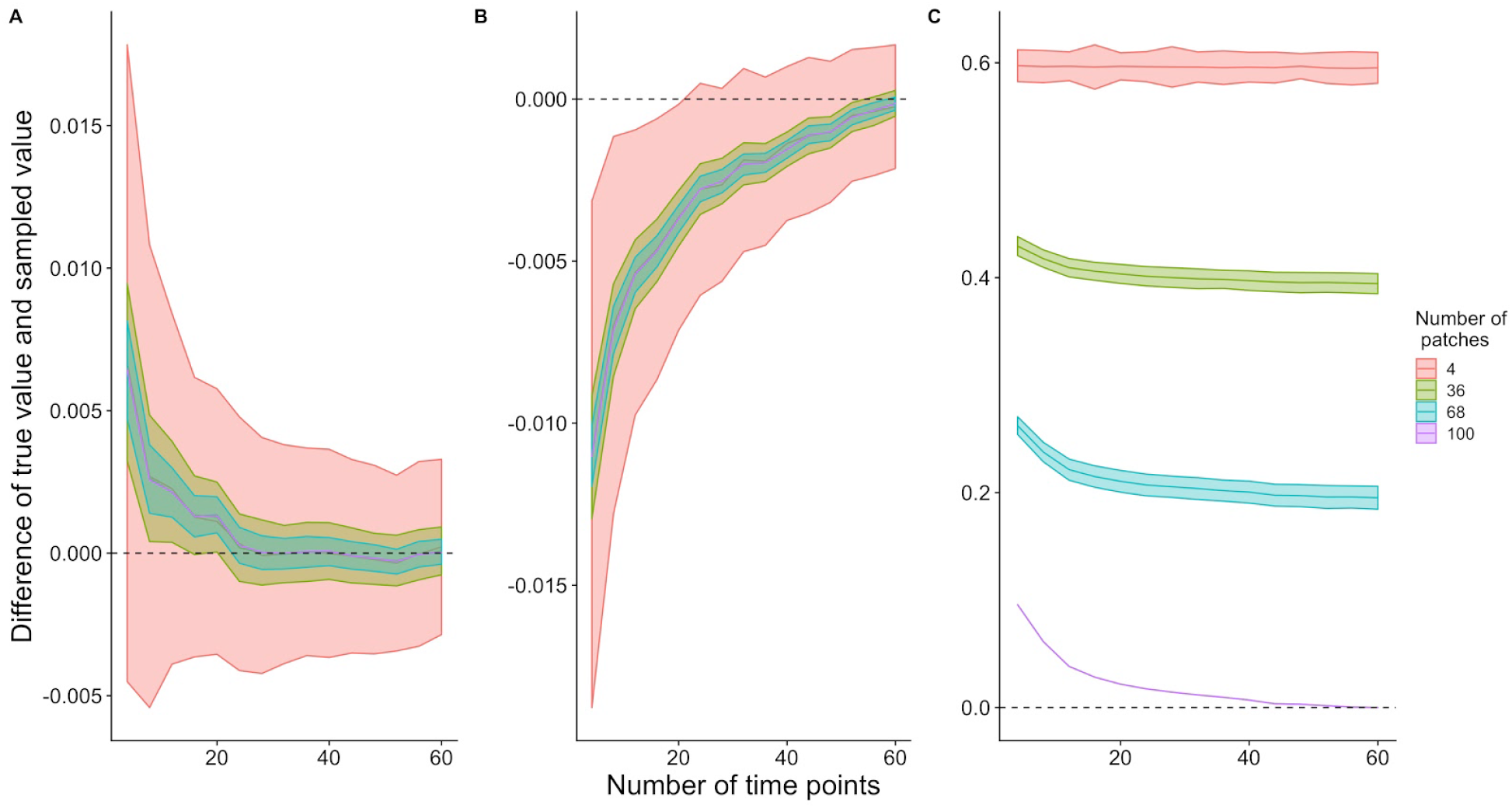
Sensitivity analysis showing how the number of time points and the number of patches sampled can increase the difference between an estimated summary statistic and its true value. Different summary statistics are more or less sensitive to the incomplete sampling of patches or time points. We present the sensitivity as the difference between the true value (i.e. the value of the summary statistic when the metacommunity was fully sampled) and the value of the summary statistic with the presented number of time points and patches. The coefficient of variation in abundance at a local scale (A) is insensitive to the number of time points or patches. The richness differences through time is very sensitive to the lack of time points (B). The mean proportion of patches occupied is more sensitive to the number of patches (C).

Some statistics are sensitive to the loss of both time points and patches. For example, the coefficient of variation in abundance at a local scale, which is critical for detecting biotic interactions, is not very sensitive to the number of time points, but the variance is higher with fewer patches sampled (Figure 4a). Richness differences through time is highly sensitive to the number of time points sampled. In addition, when the number of patches sampled is low, the variance around the summary statistics was much larger than when more patches were sampled (Figure 4b). Finally, the mean proportion of patches used by species is more sensitive to the loss of patches than time points. When the entire (100 patches) metacommunity is sampled, only 30% of the time points need to be sampled to obtain the true summary statistic. But if few patches are sampled, then even complete temporal sampling of the metacommunity will lead to large discrepancies between estimated values and true values (Figure 4c).

Multiple studies have shown that sampling effort can have large consequences for the inferences made on metacommunity processes (Gilbert and Bennett 2010, Ovaskainen et al. 2019, Viana and Chase 2019). We add to this by showing how the undersampling of the temporal series also affects the robustness of inferences. The coefficient of variation in abundance at a local scale was important for both biotic and abiotic processes. If space is fully sampled, only a few time points are needed to capture this statistic. Similarly, the environmental component of variation partitioning in space and time was a very important statistic for abiotic processes. We suggest that depending on the processes of interest, space or time may be prioritized to capture the appropriate dynamics given sampling constraints.

### General discussion and caveats

The relative importance of the summary statistics depends on the assumptions made in the simulation model. For example, in the model used here, we assumed that all the species had the same dispersal rate, though this can be changed for future studies. Interspecific variation in dispersal rates will likely reduce the degree to which observed patterns can be used to assess dispersal processes. In addition, by forcing all species to have the same dispersal rate might explain why our simulations result with empty metacommunities. We make a similar assumption for niche breadth, where all the species have the same breath but different optima and that all patches have the same size. For example, the latter assumption has consequences for the observed colonization and extinction rates (MacArthur and Wilson 1967), as well as community size and spatial environmental heterogeneity within the patch, which can weaken the relationship between dispersal rate and the summary statistics identified. By modifying different assumptions, we can identify how robust the relationship between the processes and the summary statistics will be. This approach allows us to generate hypotheses about the links between patterns and processes in metacommunities, which can then be tested via experiments and controlled observational work.

In this model we assumed that all species were governed by the same metacommunity dynamics. Empirical metacommunities may not adhere to this assumption and in fact, empirical metacommunities may have different sub-assemblages of functionally different taxa governed by different metacommunity dynamics (Thompson et al. 2017). However, this type of analysis can be useful for sets of competitors with similar dispersal abilities. In addition, we did not incorporate the complexity of trophic metacommunities (Guzman et al. 2019), but this type of workflow can be used to investigate trophic metacommunities.

While we show that the classical partitions of “E” and “S” from RDA-based variation partitioning may not be informative for inferring metacommunity processes, E and S may still be informative when we want to understand species responses to global change or to develop management actions by knowing whether the community patterns are governed more by environmental (be it indirectly through species interactions) or spatial (be it due to unmeasured environmental gradients). In addition, the comparison of S or E components between datasets or species may provide useful information (De Bie et al. 2012).

Our analysis also suggests why most empirical metacommunity analysis have such low explanatory power — dramatic undersampling in space and time. Our sensitivity analysis shows that while some of the metrics are robust to low sample size in space or time, very few are robust when both space and time are undersampled. If empirical metacommunity studies only sample few of the relevant patches or time points, trying to understand the processes that structure metacommunities might not be possible.

## Conclusions and future directions

As a next step forward, we suggest that one can use the summary statistics identified here in an Approximate Bayesian Computation (ABC) framework (Slater et al. 2012, Pontarp et al. 2019). Recent studies have also used a random forest approach to reduce dimensionality for an ABC framework (Hauenstein et al. 2019). Such a framework would use simulated models and summary statistics to determine the posterior distribution of parameters of interest (e.g.. dispersal rate) based on the distance between empirical data and summary statistics. This method can be used with an absolute model fit to assess how well the simulation model explains empirical data (Pennell et al. 2015).

Overall, we highlight two main take-home messages from our study. First, although the majority of studies of metacommunities focus on static (snapshot) patterns of species abundances and distributions in space, we showed that considering temporal dynamics is key for distinguishing the processes driving metacommunity dynamics. By doing so, our ability to explain variation in density-dependent, density-independent and dispersal processes improved by up to 60%. These results suggest that we cannot substitute space for time (or vice-versa) when we want to study metacommunity dynamics. Second, although there can never be a one-to-one matching of pattern to process, we show that it is essential to use multiple summary statistics simultaneously in order to disentangle among the fundamental processes driving metacommunity dynamics. Model based statistics in addition to descriptive statistics were needed to have the highest performance in the random forest models.

## Acknowledgements

This paper resulted from the sTURN working group funded by sDiv, the Synthesis Centre of the German Centre for Integrative Biodiversity Research (iDiv) Halle-Jena-Leipzig, funded by the German Research Foundation (FZT118). Additional funding came from Osterreichische Forschungsgemeinschaft (OFG; International Communication, project 06/15539). PLT was supported by Killam and NSERC postdoctoral fellowships. LDM acknowledges KU Leuven Research Fund project C16/2017/002 and FWO project G0B9818. ZH acknowledges support by the Interreg V-A Austria-Hungary programme of the European Regional Development Fund (project ‘Vogelwarte - Madarvarta 2’), GINOP 2.3.2.-15-2016-00057, NKFIH OTKA FK-132095, and the Janos Bolyai Research Scholarship of the Hungarian Academy of Sciences. LMG is supported by NSERC CGS-D, UBC Four Year Fellowships and SFU. JMC, DSV and AJ were also supported by the German Centre for Integrative Biodiversity Research (iDiv) Halle-Jena-Leipzig, funded by the German Research Foundation (FZT118). DSV was also supported by the Alexander von Humboldt Foundation.

