## Supplementary figures and tables for "Disentangling metacommunity processes using multiple metrics in space and time"

**Supplementary material for “Disentangling metacommunity processes using multiple metrics in space and time”**

### Appendix

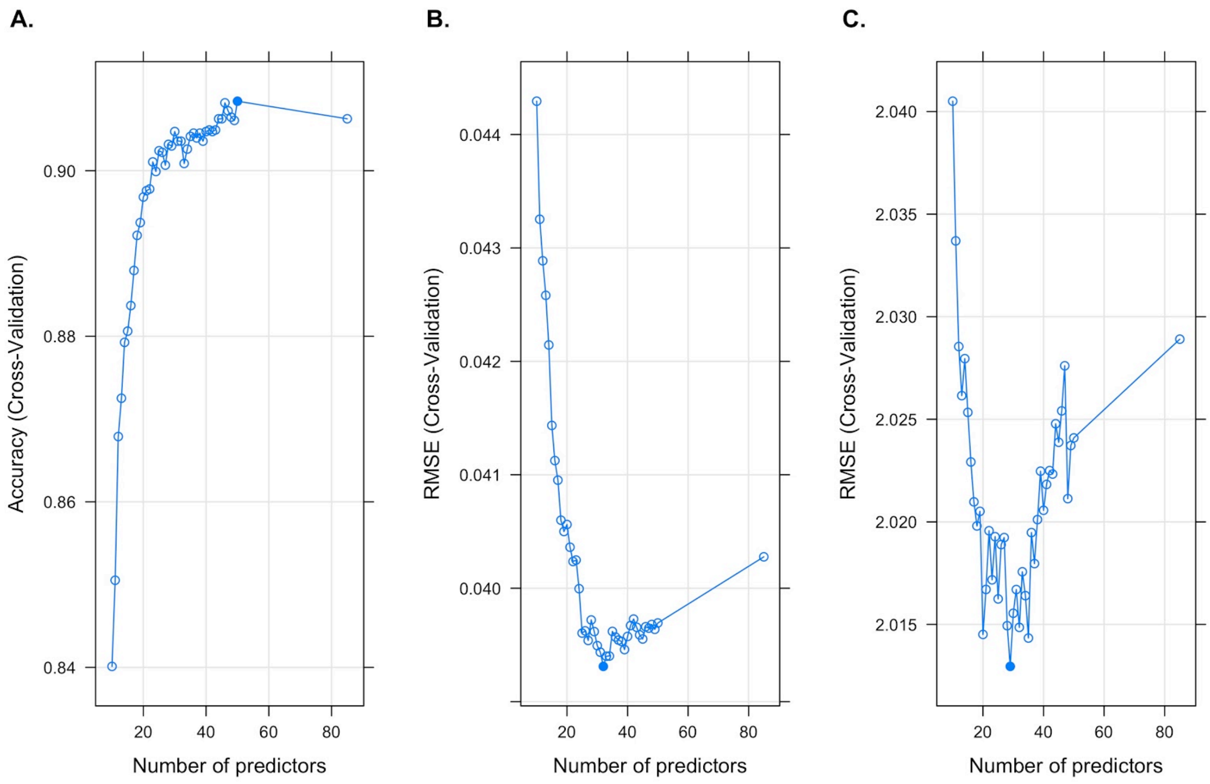


**Figure S1:** Cross validation results for the best number of predictors in the minimal model for A. density-dependent biotic interactions, B. dispersal, and C. density-independent responses to abiotic conditions. Accuracy is the proportion of times the categories were identified correctly.

**Table S1:** All of the statistics included in the analysis

| Analysis type | Description |
|---|---|
| Abundance and occupancy ratios | Ratio of abundances at local scale (Hill 1/Hill 0) |

| | |
|---|---|
| Abundance and occupancy ratios | Ratio of abundances at local scale (Hill 2/Hill 0) |
| Abundance and occupancy ratios | Ratio of occupancies at local scale (Hill 1/ Hill 0) |
| Abundance and occupancy ratios | Ratio of occupancies at local scale (Hill 2/ Hill 0) |
| Abundance and occupancy ratios | Ratio of abundances at regional scale (Hill 1/Hill 0) |
| Abundance and occupancy ratios | Ratio of abundances at regional scale (Hill 2/Hill 0) |
| CV variation | Coefficient of variation in abundance at local scale |
| CV variation | Coefficient of variation in abundance at regional scale |
| Beta diversity | Beta diversity in space (Hill 0) |
| Beta diversity | Beta diversity in space (Hill 1) |
| Beta diversity | Beta diversity in space (Hill 2) |
| Beta diversity | Beta diversity in time (Hill 0) |
| Beta diversity | Beta diversity in time (Hill 1) |
| Beta diversity | Beta diversity in time (Hill 2) |
| Beta diversity decomposition | Total beta diversity in space |
| Beta diversity decomposition | Replacement component of beta diversity in space |
| Beta diversity decomposition | Total richness difference diversity in space |
| Beta diversity decomposition | Total replacement diversity/ Total beta diversity in space |
| Beta diversity decomposition | Total richness difference diversity (or nestedness)/ Total beta diversity in space |
| Beta diversity decomposition | Total beta diversity in time |
| Beta diversity | Replacement component of beta diversity in time |

| | |
|---|---|
| decomposition | |
| Beta diversity decomposition | Total richness difference diversity in time |
| Beta diversity decomposition | Total replacement diversity/ Total beta diversity in time |
| Beta diversity decomposition | Total richness difference diversity (or nestedness)/ Total beta diversity in time |
| Proportion of patches occupied | Mean proportion of patches occupied |
| Proportion of patches occupied | Minimum proportion of patches occupied |
| Proportion of patches occupied | Maximum proportion of patches occupied |
| Variation partitioning in space | Variation partitioning in space (Space component) |
| Variation partitioning in space | Variation partitioning in space (Space and environmental component) |
| Variation partitioning in space | Variation partitioning in space (Environmental component)) |
| Variation partitioning in space and time | Variation partitioning in time and space (Space component) |
| Variation partitioning in space and time | Variation partitioning in time and space (Environmental component) |
| Variation partitioning in space and time | Variation partitioning in time and space (Time component) |
| Variation partitioning in space and time | Variation partitioning in time and space (Space and environmental component) |
| Variation partitioning in space and time | Variation partitioning in time and space (Environment and time component) |
| Variation partitioning in space and time | Variation partitioning in time and space (Space and time component) |
| Variation partitioning in space and time | Variation partitioning in time and space (Space, time and environmental component) |

| | |
|---|---|
| HMSC | HMSC mean R2 |
| HMSC | HMSC environmental component raw |
| HMSC | HMSC space-time component raw |
| HMSC | HMSC space component raw |
| HMSC | HMSC time component raw |
| HMSC | HMSC environmental component standardized |
| HMSC | HMSC space-time component standardized |
| HMSC | HMSC space component standardized |
| HMSC | HMSC time component standardized |
| HMSC | HMSC median of species associations in space-time |
| HMSC | HMSC median of species associations in space |
| HMSC | HMSC median of species associations in time |
| HMSC | HMSC mean of species associations in space-time |
| HMSC | HMSC mean of absolute species associations in space-time |
| HMSC | HMSC mean of positive species associations in space-time |
| HMSC | HMSC mean of negative species associations in space-time |
| HMSC | HMSC mean of species associations in space |
| HMSC | HMSC mean of absolute species associations in space |
| HMSC | HMSC mean of positive species associations in space |
| HMSC | HMSC mean of negative species associations in space |
| HMSC | HMSC mean of species associations in time |
| HMSC | HMSC mean of absolute species associations in time |
| HMSC | HMSC mean of positive species associations in time |
| HMSC | HMSC mean of negative species associations in time |
| HMSC | HMSC variance of species associations in space-time |
| HMSC | HMSC variance of species associations in space |

| | |
|---|---|
| HMSC | HMSC variance of species associations in time |
| HMSC | HMSC proportion of species associations higher than zero in space-time |
| HMSC | HMSC proportion of species associations lower than zero in space-time |
| HMSC | HMSC proportion of species associations equal to zero in space-time |
| HMSC | HMSC proportion of species associations higher than zero in space |
| HMSC | HMSC proportion of species associations lower than zero in space |
| HMSC | HMSC proportion of species associations equal to zero in space |
| HMSC | HMSC proportion of species associations higher than zero in time |
| HMSC | HMSC proportion of species associations lower than zero in time |
| HMSC | HMSC proportion of species associations equal to zero in space |
| HMSC | HMSC space-time POS (Proportion of species pairs with positive association above 0.95 threshold) |
| HMSC | HMSC space POS (Proportion of species pairs with positive association above 0.95 threshold) |
| HMSC | HMSC time POS (Proportion of species pairs with positive association above 0.95 threshold) |
| HMSC | HMSC space-time NEG (Proportion of species pairs with negative association below 0.05 threshold) |
| HMSC | HMSC space NEG (Proportion of species pairs with negative association below 0.05 threshold) |
| HMSC | HMSC time NEG (Proportion of species pairs with negative association below 0.05 threshold) |
| HMSC | HMSC space-time MEA (Posterior mean of spatial scale of residual variation) |

| | |
|---|---|
| HMSC | HMSC space MEA (Posterior mean of spatial scale of residual variation) |
| HMSC | HMSC time MEA (Posterior mean of spatial scale of residual variation) |
| HMSC | HMSC space-time SS (Posterior support for spatially structured residual variation) |
| HMSC | HMSC space SS (Posterior support for spatially structured residual variation) |
| HMSC | HMSC time SS (Posterior support for spatially structured residual variation) |

867

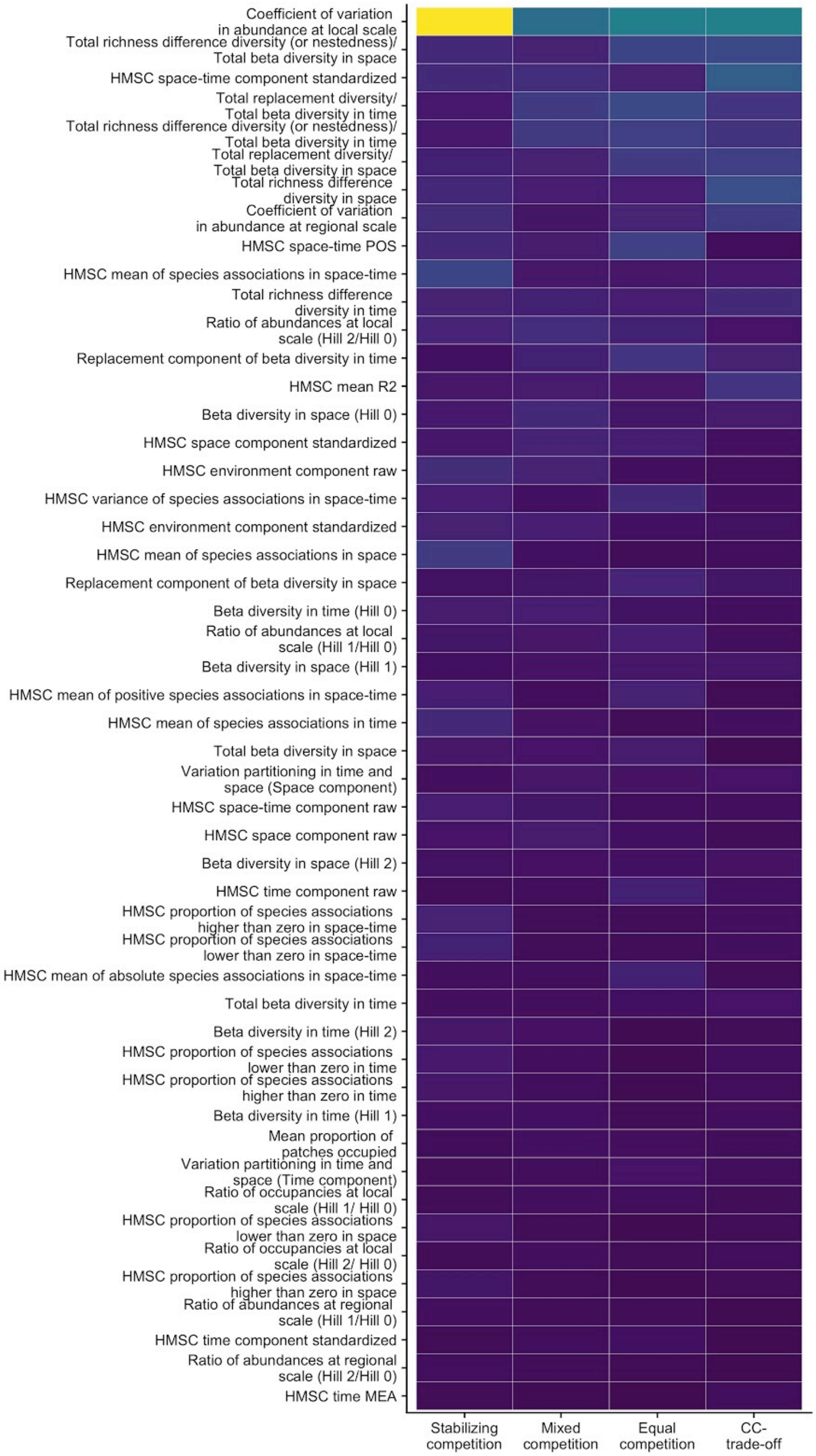

Coefficient of variation in abundance at local scale
Total richness difference diversity (or nestedness)/ Total beta diversity in space
HMSC space-time component standardized
Total replacement diversity/ Total beta diversity in time
Total richness difference diversity (or nestedness)/ Total beta diversity in time
Total replacement diversity/ Total beta diversity in space
Total richness difference diversity in space
Coefficient of variation in abundance at regional scale
HMSC space-time POS
HMSC mean of species associations in space-time
Total richness difference diversity in time
Ratio of abundances at local scale (Hill 2/Hill 0)
Replacement component of beta diversity in time
HMSC mean R2
Beta diversity in space (Hill 0)
HMSC space component standardized
HMSC environment component raw
HMSC variance of species associations in space-time
HMSC environment component standardized
HMSC mean of species associations in space
Replacement component of beta diversity in space
Beta diversity in time (Hill 0)
Ratio of abundances at local scale (Hill 1/Hill 0)
Beta diversity in space (Hill 1)
HMSC mean of positive species associations in space-time
HMSC mean of species associations in time
Total beta diversity in space
Variation partitioning in time and space (Space component)
HMSC space-time component raw
HMSC space component raw
Beta diversity in space (Hill 2)
HMSC time component raw
HMSC proportion of species associations higher than zero in space-time
HMSC proportion of species associations lower than zero in space-time
HMSC mean of absolute species associations in space-time
Total beta diversity in time
Beta diversity in time (Hill 2)
HMSC proportion of species associations lower than zero in time
HMSC proportion of species associations higher than zero in time
Beta diversity in time (Hill 1)
Mean proportion of patches occupied
Variation partitioning in time and space (Time component)
Ratio of occupancies at local scale (Hill 1/ Hill 0)
HMSC proportion of species associations lower than zero in space
Ratio of occupancies at local scale (Hill 2/ Hill 0)
HMSC proportion of species associations higher than zero in space
Ratio of abundances at regional scale (Hill 1/Hill 0)
HMSC time component standardized
Ratio of abundances at regional scale (Hill 2/Hill 0)
HMSC time MEA
Stabilizing competition
Mixed competition
Equal competition
CC-trade-off

**Figure S2:** Importance of the predictors in the minimal model for the density-dependent biotic interactions process. Ordered from more important to less important similar to Figure 2.

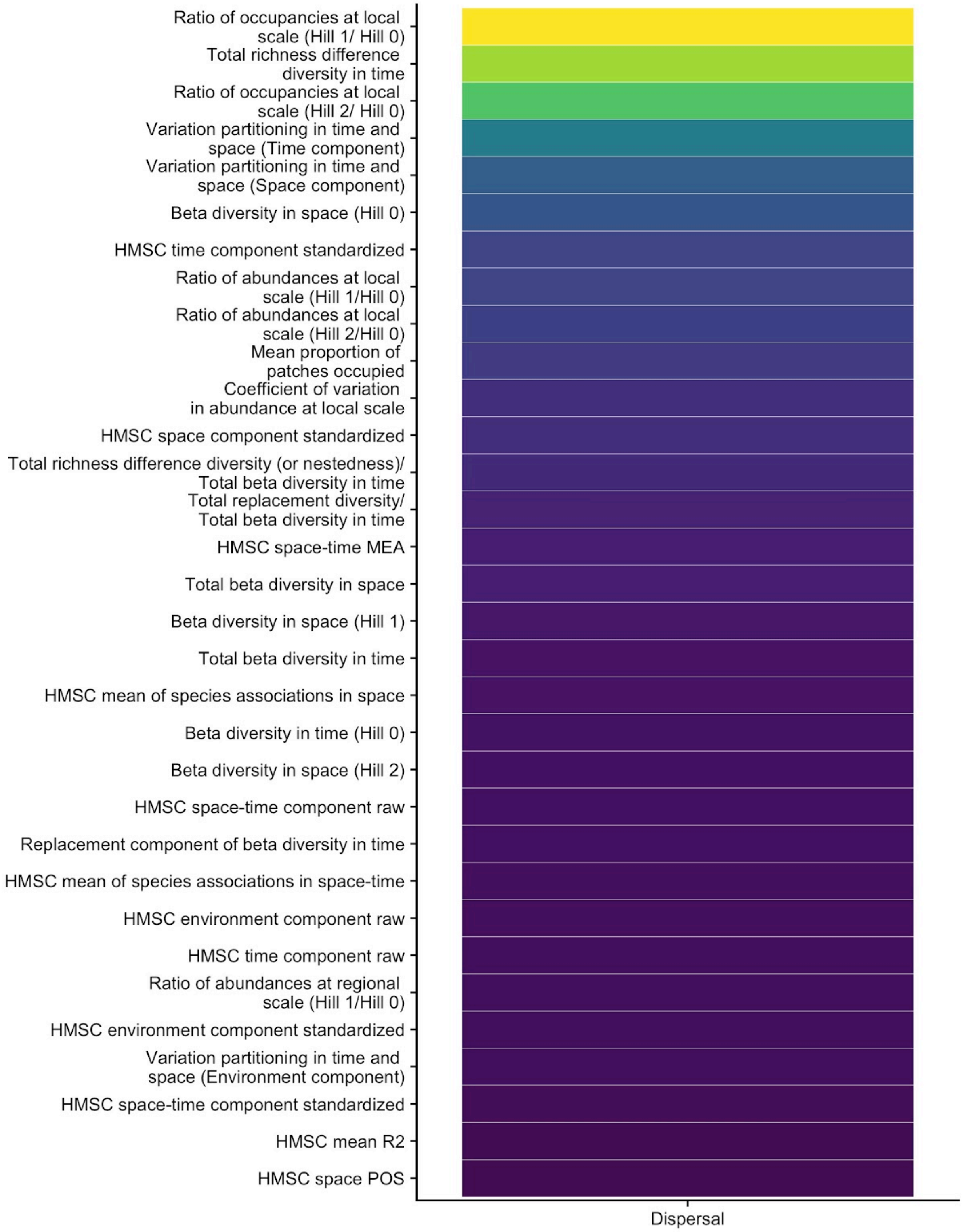
Ratio of occupancies at local scale (Hill 1/ Hill 0)
Total richness difference diversity in time
Ratio of occupancies at local scale (Hill 2/ Hill 0)
Variation partitioning in time and space (Time component)
Variation partitioning in time and space (Space component)
Beta diversity in space (Hill 0)
HMSC time component standardized
Ratio of abundances at local scale (Hill 1/Hill 0)
Ratio of abundances at local scale (Hill 2/Hill 0)
Mean proportion of patches occupied
Coefficient of variation in abundance at local scale
HMSC space component standardized
Total richness difference diversity (or nestedness)/ Total beta diversity in time
Total replacement diversity/ Total beta diversity in time
HMSC space-time MEA
Total beta diversity in space
Beta diversity in space (Hill 1)
Total beta diversity in time
HMSC mean of species associations in space
Beta diversity in time (Hill 0)
Beta diversity in space (Hill 2)
HMSC space-time component raw
Replacement component of beta diversity in time
HMSC mean of species associations in space-time
HMSC environment component raw
HMSC time component raw
Ratio of abundances at regional scale (Hill 1/Hill 0)
HMSC environment component standardized
Variation partitioning in time and space (Environment component)
HMSC space-time component standardized
HMSC mean R2
HMSC space POS
Dispersal

**Figure S3:** Importance of the predictors in the minimal model for the dispersal process. Ordered from more important to less important similar to Figure 2.

875

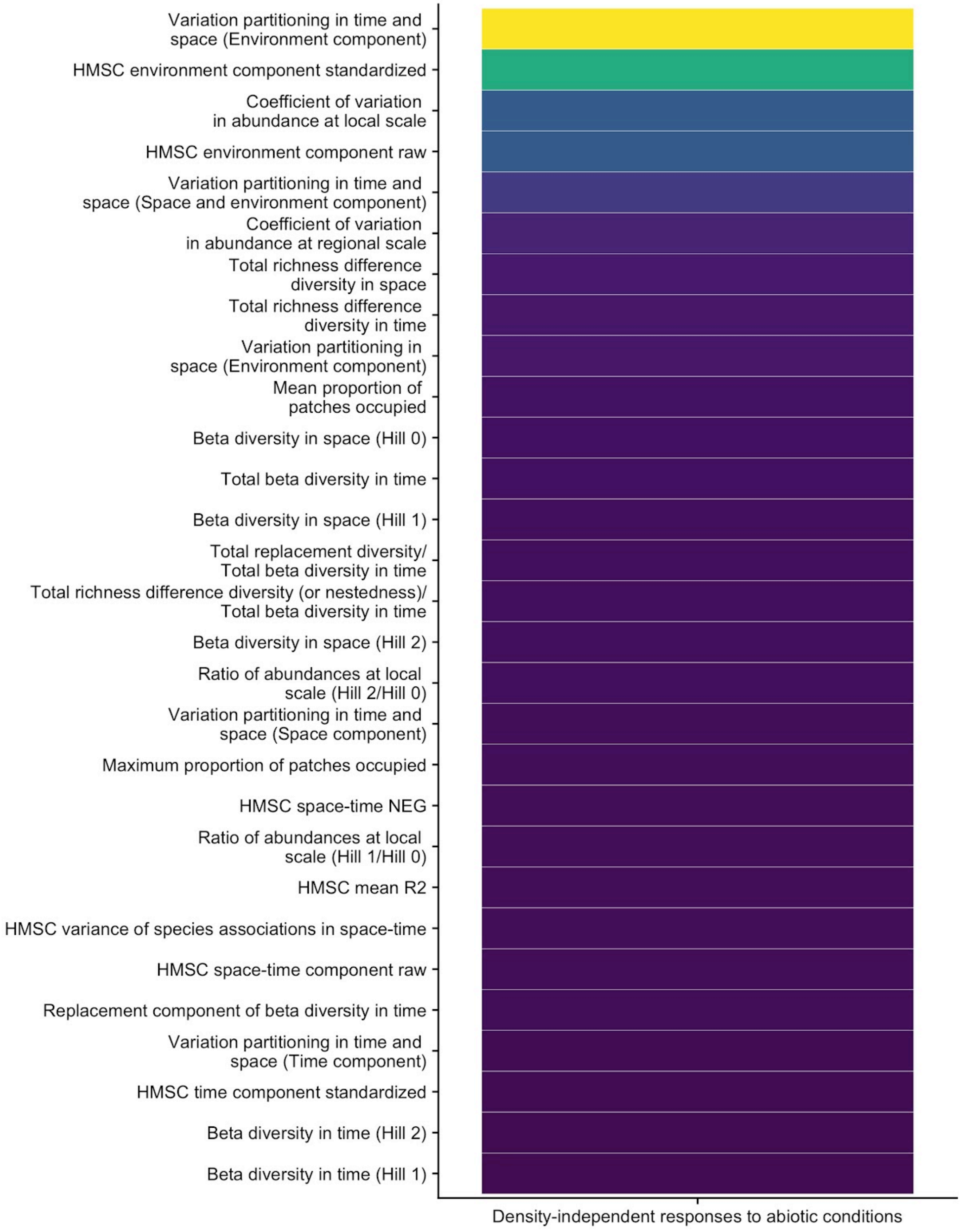
Variation partitioning in time and space (Environment component)
HMSC environment component standardized
Coefficient of variation in abundance at local scale
HMSC environment component raw
Variation partitioning in time and space (Space and environment component)
Coefficient of variation in abundance at regional scale
Total richness difference diversity in space
Total richness difference diversity in time
Variation partitioning in space (Environment component)
Mean proportion of patches occupied
Beta diversity in space (Hill 0)
Total beta diversity in time
Beta diversity in space (Hill 1)
Total replacement diversity/ Total beta diversity in time
Total richness difference diversity (or nestedness)/ Total beta diversity in time
Beta diversity in space (Hill 2)
Ratio of abundances at local scale (Hill 2/Hill 0)
Variation partitioning in time and space (Space component)
Maximum proportion of patches occupied
HMSC space-time NEG
Ratio of abundances at local scale (Hill 1/Hill 0)
HMSC mean R2
HMSC variance of species associations in space-time
HMSC space-time component raw
Replacement component of beta diversity in time
Variation partitioning in time and space (Time component)
HMSC time component standardized
Beta diversity in time (Hill 2)
Beta diversity in time (Hill 1)
Density-independent responses to abiotic conditions

**Figure S4:** Importance of the predictors in the minimal model for the density-independent responses to abiotic conditions. Ordered from more important to less important similar to Figure 2.

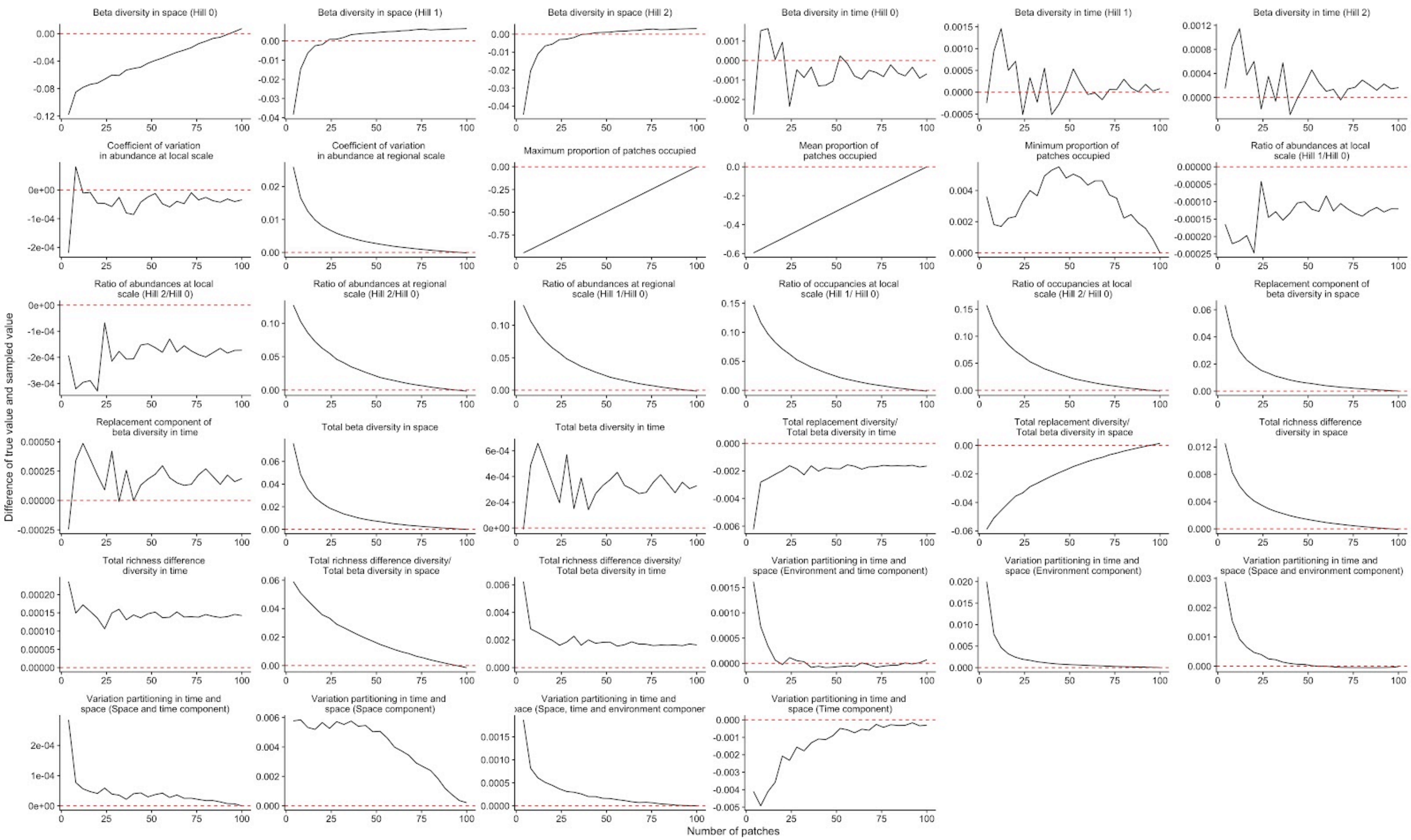
Beta diversity in space (Hill 0)
Beta diversity in space (Hill 1)
Beta diversity in space (Hill 2)
Beta diversity in time (Hill 0)
Beta diversity in time (Hill 1)
Beta diversity in time (Hill 2)
Coefficient of variation in abundance at local scale
Coefficient of variation in abundance at regional scale
Maximum proportion of patches occupied
Mean proportion of patches occupied
Minimum proportion of patches occupied
Ratio of abundances at local scale (Hill 1/Hill 0)
Ratio of abundances at local scale (Hill 2/Hill 0)
Ratio of abundances at regional scale (Hill 2/Hill 0)
Ratio of abundances at regional scale (Hill 1/Hill 0)
Ratio of occupancies at local scale (Hill 1/ Hill 0)
Ratio of occupancies at local scale (Hill 2/ Hill 0)
Replacement component of beta diversity in space
Replacement component of beta diversity in time
Total beta diversity in space
Total beta diversity in time
Total replacement diversity/ Total beta diversity in time
Total replacement diversity/ Total beta diversity in space
Total richness difference diversity in space
Total richness difference diversity in time
Total richness difference diversity/ Total beta diversity in space
Total richness difference diversity/ Total beta diversity in time
Variation partitioning in time and space (Environment and time component)
Variation partitioning in time and space (Environment component)
Variation partitioning in time and space (Space and environment component)
Variation partitioning in time and space (Space and time component)
Variation partitioning in time and space (Space component)
Variation partitioning in time and space (Space, time and environment componer
Variation partitioning in time and space (Time component)
Difference of true value and sampled value
Number of patches

**Figure S5:** Difference between the estimated value and the true value when the metacommunity is fully sampled temporally (t = 60) when reducing the number of sampled patches.

**Table S2:** "Half life" in patches of each of the summary statistics. The half life is the minimum number of patches (from a total of 100) needed to reduce the 'error' in the summary statistic by half.

| **Statistic** | **Half life in patches** |
|---|---|
| Total beta diversity in time | 4 |
| Beta diversity in space (Hill 1) | 8 |
| Beta diversity in space (Hill 2) | 8 |
| Minimum proportion of patches occupied | 8 |
| Total replacement diversity/ Total beta diversity in time | 8 |
| Total richness difference diversity/ Total beta diversity in time | 8 |
| Variation partitioning in time and space (Environmental component) | 8 |
| Variation partitioning in time and space (Space and environmental component) | 8 |
| Variation partitioning in time and space (Environment and time component) | 8 |

| | |
|---|---|
| Variation partitioning in time and space (Space and time component) | 8 |
| Variation partitioning in time and space (Space, time and environmental component) | 8 |
| Coefficient of variation in abundance at regional scale | 12 |
| Total beta diversity in space | 12 |
| Replacement component of beta diversity in space | 12 |
| Total richness difference diversity in space | 12 |
| Ratio of occupancies at local scale (Hill 2/ Hill 0) | 16 |
| Ratio of occupancies at local scale (Hill 1/ Hill 0) | 20 |
| Ratio of abundances at regional scale (Hill 1/Hill 0) | 20 |
| Ratio of abundances at regional scale (Hill 2/Hill 0) | 20 |
| Variation partitioning in time and space (Time component) | 20 |
| Total richness difference diversity in time | 24 |
| Ratio of abundances at local scale (Hill 2/Hill 0) | 24 |

| | |
|---|---|
| Beta diversity in space (Hill 0) | 28 |
| Total replacement diversity/ Total beta diversity in space | 28 |
| Total richness difference diversity/ Total beta diversity in space | 28 |
| Replacement component of beta diversity in time | 32 |
| Beta diversity in time (Hill 0) | 40 |
| Coefficient of variation in abundance at local scale | 40 |
| Mean proportion of patches occupied | 52 |
| Maximum proportion of patches occupied | 52 |
| Beta diversity in time (Hill 2) | 60 |
| Ratio of abundances at local scale (Hill 1/Hill 0) | 60 |
| Beta diversity in time (Hill 1) | 68 |
| Variation partitioning in time and space (Space component) | 72 |

938

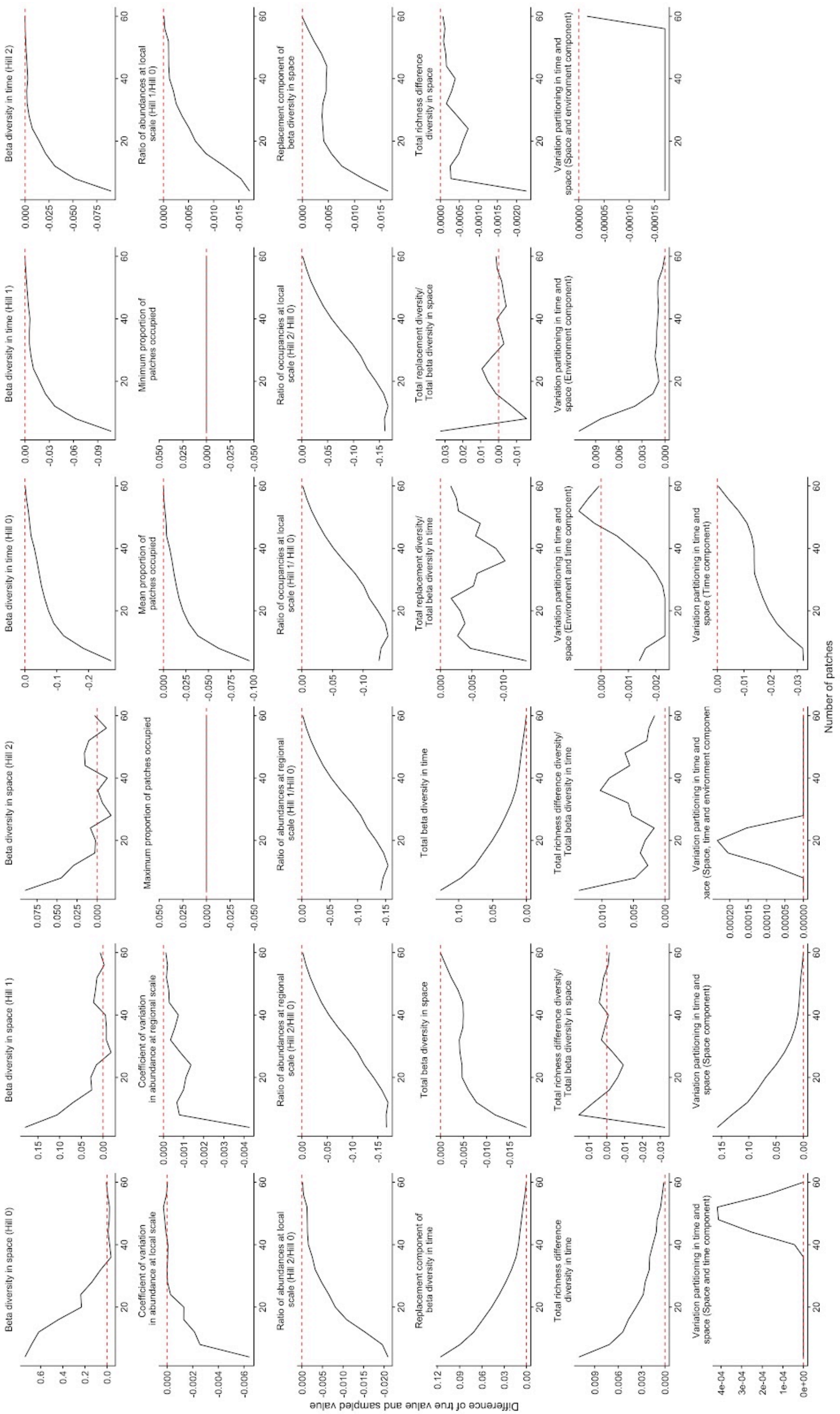
Beta diversity in space (Hill 0)
Beta diversity in space (Hill 1)
Beta diversity in space (Hill 2)
Beta diversity in time (Hill 0)
Beta diversity in time (Hill 1)
Beta diversity in time (Hill 2)
Coefficient of variation in abundance at local scale
Coefficient of variation in abundance at regional scale
Maximum proportion of patches occupied
Mean proportion of patches occupied
Minimum proportion of patches occupied
Ratio of abundances at local scale (Hill 1/Hill 0)
Ratio of abundances at local scale (Hill 2/Hill 0)
Ratio of abundances at regional scale (Hill 2/Hill 0)
Ratio of abundances at regional scale (Hill 1/Hill 0)
Ratio of occupancies at local scale (Hill 1/ Hill 0)
Ratio of occupancies at local scale (Hill 2/ Hill 0)
Replacement component of beta diversity in space
Replacement component of beta diversity in time
Total beta diversity in space
Total beta diversity in time
Total replacement diversity/ Total beta diversity in time
Total replacement diversity/ Total beta diversity in space
Total richness difference diversity in space
Total richness difference diversity in time
Total richness difference diversity/ Total beta diversity in space
Total richness difference diversity/ Total beta diversity in time
Variation partitioning in time and space (Environment and time component)
Variation partitioning in time and space (Environment component)
Variation partitioning in time and space (Space and environment component)
Variation partitioning in time and space (Space and time component)
Variation partitioning in time and space (Space component)
Variation partitioning in time and space (Time component)
Difference of true value and sampled value
Number of patches

**Figure S6:** Difference between the estimated value and the true value when the metacommunity is fully sampled spatially (patches = 100) when reducing the number of time points sampled.

**Table S3:** "Half life" in timepoints of each of the summary statistics. The half life is the minimum number of time points (from a total of 60) needed to reduce the 'error' in the summary statistic by half.

| **Statistic** | **Half life in time points** |
|---|---|
| Minimum proportion of patches occupied | 4 |
| Maximum proportion of patches occupied | 4 |
| Variation partitioning in time and space (Space and time component) | 4 |
| Variation partitioning in time and space (Space, time and environmental component) | 4 |
| Beta diversity in space (Hill 1) | 8 |
| Beta diversity in space (Hill 2) | 8 |
| Beta diversity in time (Hill 1) | 8 |
| Beta diversity in time (Hill 2) | 8 |
| Coefficient of variation in abundance at local scale | 8 |

| | |
|---|---|
| Beta diversity in time (Hill 0) | 12 |
| Mean proportion of patches occupied | 12 |
| Total beta diversity in space | 12 |
| Replacement component of beta diversity in space | 12 |
| Total richness difference diversity in time | 12 |
| Variation partitioning in time and space (Environmental component) | 12 |
| Beta diversity in space (Hill 0) | 16 |
| Total beta diversity in time | 16 |
| Replacement component of beta diversity in time | 16 |
| Ratio of abundances at local scale (Hill 1/Hill 0) | 16 |
| Ratio of abundances at local scale (Hill 2/Hill 0) | 16 |
| Variation partitioning in time and space (Space component) | 16 |
| Coefficient of variation in abundance at regional scale | 24 |

| | |
|---|---|
| Total richness difference diversity in space | 24 |
| Total replacement diversity/ Total beta diversity in space | 24 |
| Total richness difference diversity/ Total beta diversity in space | 24 |
| Variation partitioning in time and space (Time component) | 28 |
| Ratio of abundances at regional scale (Hill 2/Hill 0) | 32 |
| Ratio of occupancies at local scale (Hill 1/ Hill 0) | 36 |
| Ratio of occupancies at local scale (Hill 2/ Hill 0) | 36 |
| Ratio of abundances at regional scale (Hill 1/Hill 0) | 36 |
| Variation partitioning in time and space (Environment and time component) | 44 |
| Total replacement diversity/ Total beta diversity in time | 48 |
| Total richness difference diversity/ Total beta diversity in time | 48 |
| Variation partitioning in time and space (Space and environmental component) | 60 |

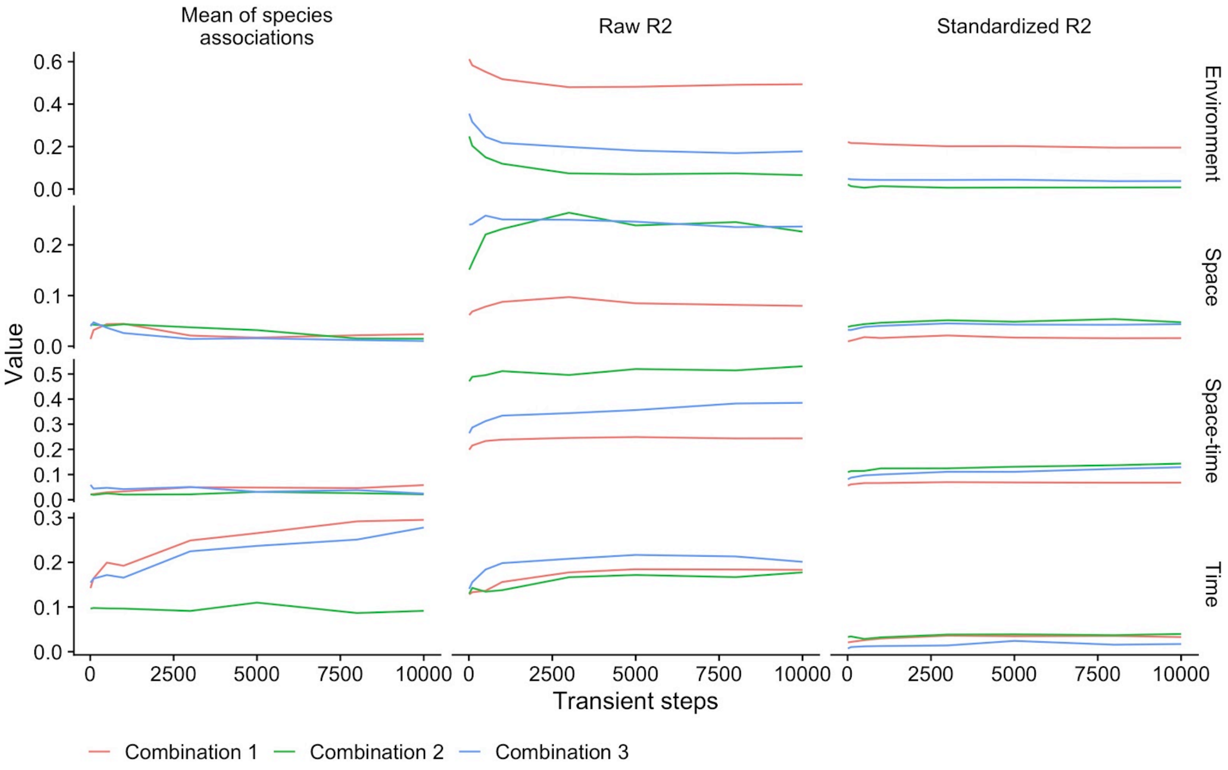


**Figure S7:** Sensitivity analysis to assess whether 5000 time steps is sufficient for estimating the summary statistics from the HMSC models. We performed the sensitivity analysis on three metrics: the raw R2, the standardized R2, and the mean species association for the four main predictors. We also did the sensitivity analysis on three combinations of dispersal, density-independent abiotic niche, and density-dependent species interaction values. For combination 1 we used dispersal ($a_i$) = 0.001, the abiotic niche breadth parameter ($\sigma_i$) = 0.46416, and equal competition. For combination 2, $a_i$ = 0.001, $\sigma_i$ = 0.46416 and stabilizing competition. For combination 3, $a_i$ = 0.001, $\sigma_i$ = 10 and stabilizing competition. All sensitivity analyses were based on a single replicate run of the simulated data for each parameter combination. This analysis shows that the HMSC model run with 5000 time steps, as we used in our main analysis, provides reasonable estimates of the summary statistics. That is, that the estimates differ among scenarios in our simulation model, and that running the HMSC model with more time steps does not qualitatively change the estimated summary statistics.
